# Mechanical and Interfacial Coupling in Bacterial Biofilm Detachment Underwater

**DOI:** 10.64898/2026.11.20.762633

**Authors:** Amado Luis K. Orden, Aaron Joseph Brennan, Luz Amelia Calderón-Vergara, Sumeet Kumar Das, Muskan, Amal Narayanan

**Affiliations:** George and Josephine Butler Polymer Research Laboratory, Center for Macromolecular Science and Engineering, Department of Chemistry, University of Florida, Gainesville, Florida 32611, United States

## Abstract

Biofilms form persistent surface-associated communities across natural, industrial, and biomedical environments and cause significant socioeconomic distress, yet their controlled removal remains challenging, especially in wet and underwater conditions. Inspired by barnacle adhesives that clear underlying biomass before their permanent attachment, here we introduce “adhesive peeling” as a readily adoptable, practical strategy for controlled removal and characterization of biofilms underwater. Using underwater adhesives, force-controlled contact peeling, and *in situ* confocal microscopy, we show that bacterial biofilms can be transferred from submerged surfaces to an adhesive, with removal reaching ∼ 82%. Additionally, with a mechano-interfacial map that combines underwater biofilm–adhesive work of adhesion and biofilm–adhesive mechanical mismatch, we found three distinct species-dependent adhesive peeling mechanisms: (i) *E. coli* peeling is dominated by favorable interfacial interactions and is insensitive to adhesive mechanics, (ii) *S. aureus* peeling requires a balance between interfacial affinity and mechanical response, and (iii) *P. aeruginosa* peeling is insensitive to adhesive identity, consistent with cohesion-limited failure within the biofilm. These results establish adhesive peeling as a methodology for both removing biofilms and identifying the dominant failure pathway within submerged adhesive-biofilm-substrate-water interfaces, providing a physical basis for understanding how living biofilms contact, adhere, and fail against non-living materials across healthcare, transportation, and industrial systems where persistent biofouling carries substantial human and economic costs.

## 1.0 INTRODUCTION

Bacteria predominantly exist in nature as surface-associated multicellular communities embedded within a self-produced extracellular matrix known as biofilms.^1–3^ The living and non-living constituents of this matrix collectively impart emergent properties, including interfacial adhesion, mechanical robustness, and long-term persistence, making biofilms one of the most prevalent living structures on Earth.^4,5^ Biofilms colonize nearly every wetted surface, including natural, biological, industrial, and biomedical surfaces ranging from rocks and ships to medical implants and host tissues.^6–8^ While biofilms provide important ecological and biological benefits to humanity,^9–11^ their persistent attachment to surfaces also contributes to a major economic burden worldwide through chronic infections, biofouling, and material degradation.^12–14^

Current strategies for managing undesirable biofilms include antimicrobial agents, enzymatic degradation of the extracellular matrix, antifouling surfaces and coatings, and mechanical abrasion.^15–22^ Despite these considerable advances, the removal of biofilms remains remarkably challenging, especially in submerged environments, where an intervening aqueous phase separates biofilms from the contacting treatment. Consequently, many antibiofilm strategies only reduce microbial viability or prevent initial bacterial adhesion without effectively removing the adherent biomass from the underlying substrate.^23–28^

While existing antibiofilm strategies are largely inefficient against submerged biofilms, marine macrofoulers such as barnacles and mussels routinely establish permanent attachment in environments where surfaces are frequently conditioned by microbial communities.^29–32^ For example, barnacles actively reorganize, displace, and condition underlying biological material at the interface prior to their permanent attachment, indicating that their underwater adhesion requires the removal of underlying biofilms.^33,34^ These observations, routine in nature, led us to hypothesize that synthetic underwater adhesives could similarly compete with biofilm–substrate interactions and initiate detachment through a controlled adhesive peeling process (Figure 1A).

**Figure 1.**
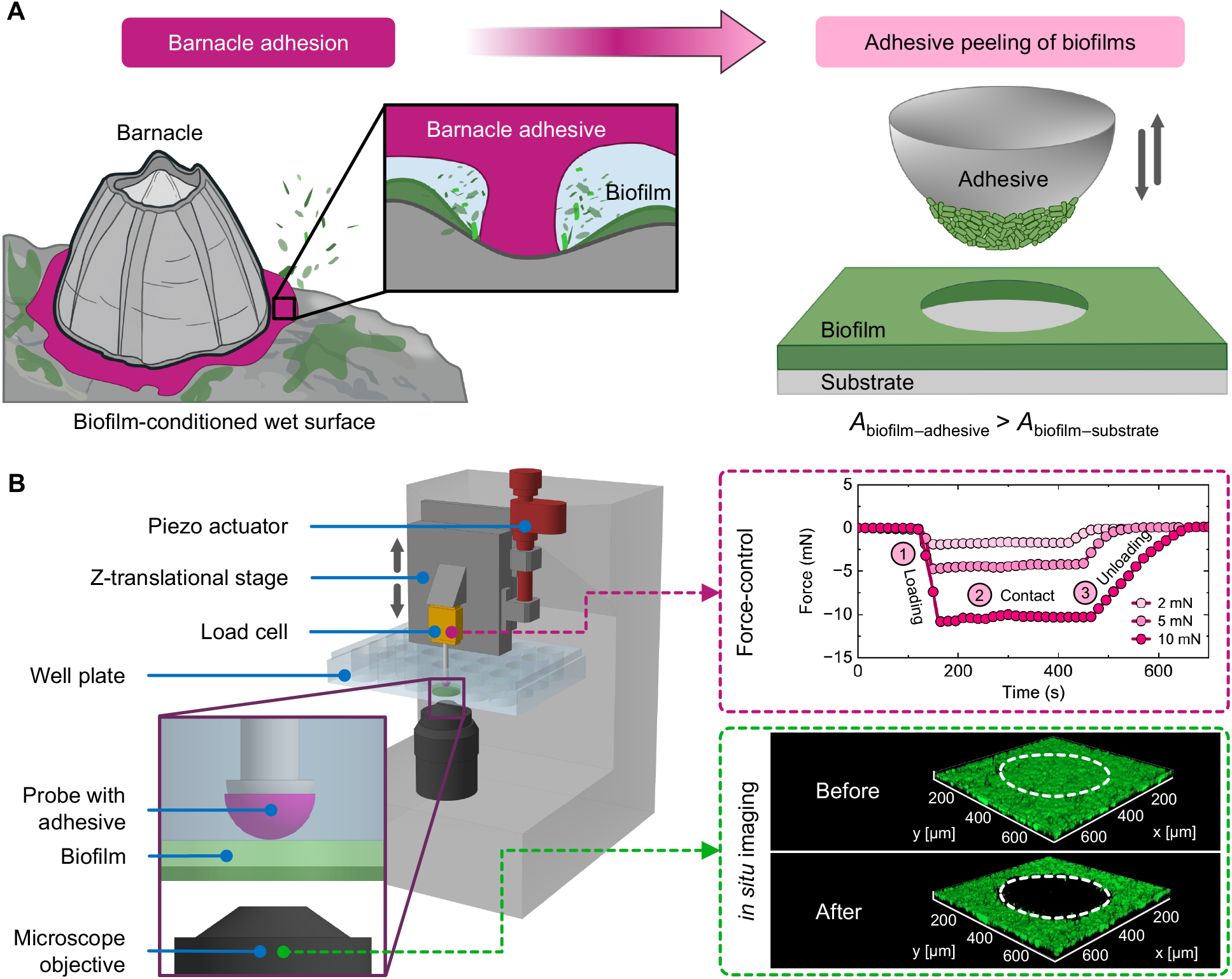
Bioinspired strategy and experimental platform for underwater biofilm peeling. (A) Barnacles displace biofilms underwater, inspiring the use of polydimethylsiloxane (PDMS) as an underwater biofilm peeler adhesive. We hypothesize that successful peeling and transfer occur when the adhesion between the biofilm and the adhesive is greater than the adhesion between the biofilm and the substrate. (B) The experimental platform using controlled biofilm peeling conditions shows the force response profile and *in situ* visualization. The force profile shows representative force curves during biofilm peeling, corresponding to 2, 5, and 10 mN contact loading, fixed 300 s contact, and fixed peeling rate. Biofilm peeling was quantified from the differences in biofilm fluorescence intensity in the contact area (dashed white outlines) before (top) and after (bottom) peeling.

Inspired by marine macrofoulers’ strategies for underwater adhesive priming, in this study, we introduce “adhesive peeling of biofilms” as a practical approach for removing biofilms underwater at scale and investigating how the peeling response reflects the material properties of the detaching interfaces. Using a family of synthetic underwater adhesives spanning a broad range of adhesive and cohesive properties, a force-controlled peeling platform, and *in situ* confocal microscopy, we systematically examined the underwater detachment of biofilms across multiple bacterial species and substrate chemistries under controlled loading conditions. We identify distinct peeling regimes associated with adhesive failure, cohesive failure, and successful biofilm transfer, and demonstrate that the outcome of peeling is governed by the interplay between adhesive interactions, cohesive material properties, interfacial chemistry, and biofilm identity. Notably, 25 µg of our adhesive removes approximately 6 nL of established *Staphylococcus aureus* (*S. aureus*) and *Escherichia coli* (*E. coli*) biofilms over a 0.25 mm² contact area, corresponding to a treatment capacity of approximately 100 cm^2^ per gram of adhesive. These results establish adhesive peeling as a mechano-interfacial framework for studying species- and substrate-dependent biofilm detachment and provide design principles for the removal of persistent biofilms from submerged surfaces.

## 2.0 RESULTS

### 2.1 Adhesive peeling enables the detachment of biofilms from substrates underwater

We hypothesized that biofilm removal could be achieved through a competitive adhesive process in which a sufficiently adhesive material transfers the biofilm from the underlying substrate to the contacting surface (Figure 1A).^32,34,35^ In this framework, when adhesion between the biofilm and adhesive polymer, *A*_biofilm–adhesive,_ is greater than adhesion between the biofilm and substrate, *A*_biofilm–substrate_, the biofilm will transfer to the adhesive. We refer to this process as “adhesive peeling of biofilms” (Figure 1A). The design of an adhesive biofilm peeler with *A*_biofilm–adhesive_ > *A*_biofilm–substrate_ underwater was guided by an established underwater contact mechanics framework, where the interfacial contact with a hydrophobic surface is energetically favored underwater, leading to molecular contact and adhesive interactions.^36–39^ Based on this framework, we reasoned that a deformable hydrophobic elastomer may establish true molecular contact with biofilms and promote their transfer during peeling underwater. We selected a commercial hydrophobic elastomer, polydimethylsiloxane (PDMS, Sylgard 184), and a family of fluorescent hydrophobic polyesters (HPE) with tunable mechanical properties for establishing the adhesive peeling of biofilms.^37,40,41^

First, we developed a force-controlled peeling method that combines predetermined loading, controlled contact time, and *in situ* fluorescent confocal microscopy (Figure 1B). In the preliminary experiment, PDMS was brought into contact with a bacterial biofilm under a contact load of 10 mN, equilibrated for 300 s, and unloaded at a rate of 30 nm s^-1^. The representative force profile consisted of an approach and loading phase, force-controlled contact equilibration, and unloading (Figure 1B). Upon unloading, we observed that *S. aureus* biofilms could be detached from submerged glass surfaces using PDMS through adhesive peeling (Figure 1B). The contact interface imaging using confocal microscopy revealed a loss of ∼ 81.6 ± 8.7 % biomass from a circular area of peel of ∼ 0.25 mm^2^. The controlled loss of biomass from this defined circular contact area demonstrates that biofilms can be mechanically removed through competitive adhesive interactions underwater. These preliminary observations establish adhesive peeling as a mechanism for underwater biofilm detachment and directed us to explore how adhesive interactions, material properties, substrate chemistry, and biofilm identity govern the adhesive peeling of biofilms underwater.

### 2.2 Hydrophobic adhesives can peel *S. aureus* biofilms from glass

The adhesive peeling framework (Figure 1A) predicts that biofilm removal should increase with increasing adhesive contact area and interactions at the biofilm–adhesive interface.^42,43^ To test this hypothesis, we compared the water contact angles (WCAs) of hydrophobic PDMS (109.9 ± 1.5°) with surface-oxidized PDMS (ox-PDMS), which possesses identical mechanical properties to those of PDMS, but shows substantially lower hydrophobicity (26.0 ± 6.3°) (Figure 2A). The contact mechanics framework predicts weaker biofilm–adhesive interactions for biofilm–ox-PDMS compared with biofilm–PDMS.^37,39,44^

**Figure 2.**
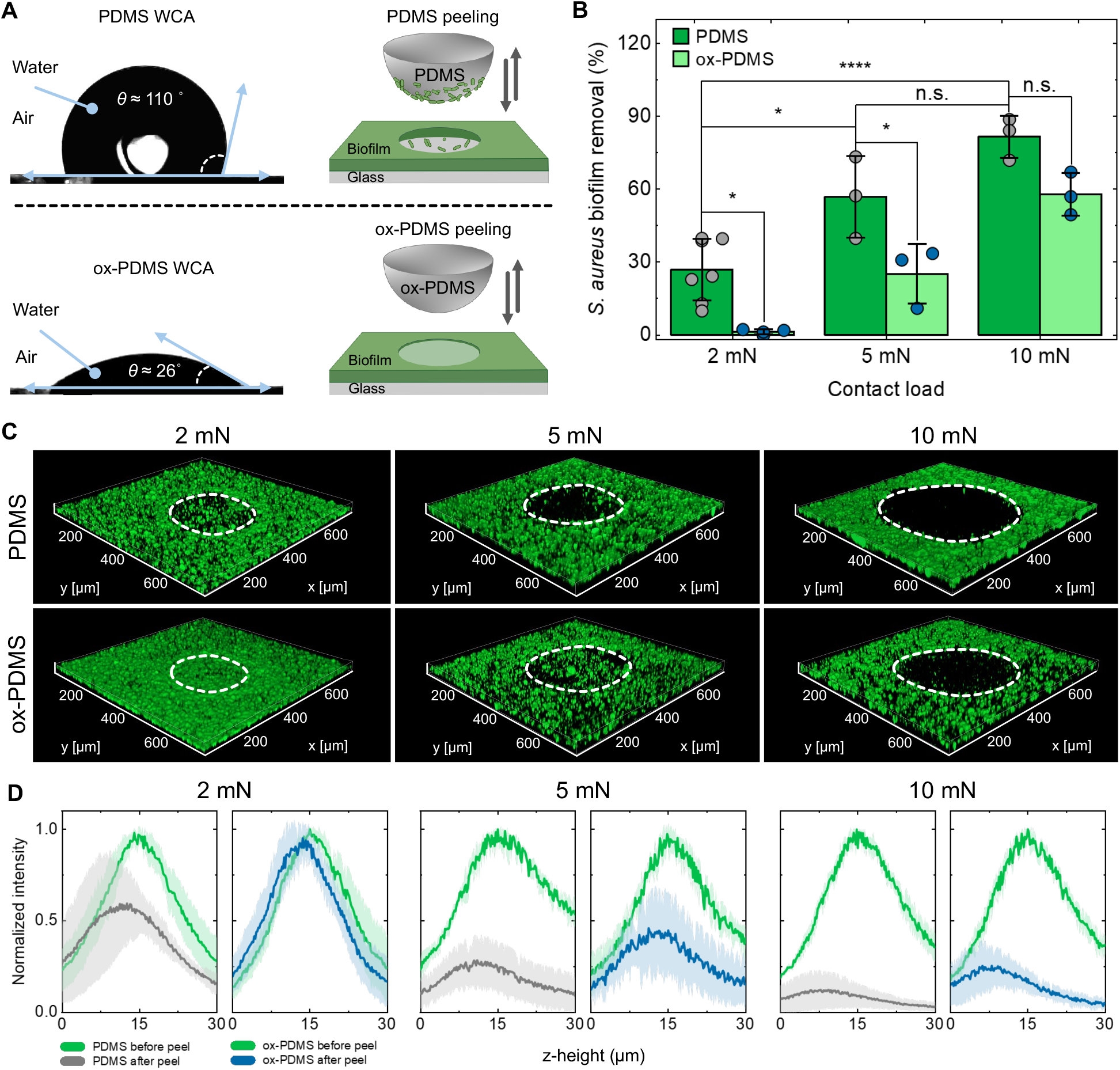
Effects of interface chemistry and contact load on the adhesive peeling of *S. aureus* biofilms underwater. (A) The water contact angle (WCA) of PDMS and surface-oxidized PDMS (ox-PDMS) and their schematic relation to biofilm peeling ability. Because of its hydrophobicity, PDMS is expected to promote water displacement and form interfacial contact with the biofilm underwater, whereas ox-PDMS is expected to retain interfacial water and limit biofilm contact and peeling. (B) *S. aureus* biofilm removal by PDMS (gray symbol) and ox-PDMS (blue symbol) at contact loads of 2, 5, and 10 mN. Bar height represents the mean, error bars show standard deviation, and the circular symbols show individual replicates. PDMS removed significantly more *S. aureus* biofilm than ox-PDMS at 2 and 5 mN; the difference at 10 mN was not statistically significant. Mean comparisons were performed using one-way ANOVA, and the significant differences across individual conditions were determined using Tukey’s test (n.s. = not significant, \**p* < 0.05, \*\**p* < 0.01, \*\*\**p* <0.001, \*\*\*\**p* <0.0001). (C) Representative Z-stack images after peeling with PDMS and ox-PDMS at 2, 5, and 10 mN contact load. Dashed white outlines indicate the contact region, which becomes progressively clearer as load is increased. (D) Normalized fluorescence intensity before and after peeling as a function of depth through the *S. aureus* thickness within the contact region at 2, 5, and 10 mN contact loads. The reduction in signal is observed at all depths, with greater reductions observed at higher contact loads. PDMS showed a greater reduction in fluorescence intensity than ox-PDMS at lower contact loads and throughout the biofilm thickness.

*S. aureus* biofilms grown on a glass surface were used as a model system and subjected to adhesive peeling at contact loads of 2, 5, and 10 mN. Z-stack images were generated for each contact load of PDMS and ox-PDMS before and after peeling, allowing visualization of biofilm removal. The corresponding intensity plots for both materials showed increasing biofilm removal with increasing contact load. PDMS removed 26.9 ± 12.8 %, 56.8 ± 16.8 %, and 81.6 ± 8.7 % of the biofilms at contact loads of 2, 5, and 10 mN, respectively (Figure 2B). In contrast, ox-PDMS removed only 1.3 ± 1.1 %, 25.1 ± 12.3 %, and 57.8 ± 8.8 % under identical conditions (Figure 2B). The analysis of fluorescence intensity across the Z-direction showed biomass loss throughout the biofilm thickness, with progressively greater reduction at higher contact loads (Figure 2C and Figure 2D). The spatial heterogeneity in biofilm removal% within the x-y contact area plane is consistent with patchy underwater contact between the hydrophobic adhesive and the hydrated biofilm (Figure 2D), in which regions of direct molecular contact coexist with regions containing trapped interfacial water. Such patchy contact is characteristic of hydrophobic–hydrophilic interfacial contact underwater and limits peeling to regions where interfacial water is sufficiently displaced to establish biofilm–adhesive contact, preventing clean ∼ 100% removal of the *S. aureus* biofilms.^38,39,41^

The consistently higher removal efficiency observed for PDMS supports the hypothesis that the energetically favored hydrophobic underwater contact contributes to underwater biofilm peeling. The greater removal achieved at lower contact loads with PDMS suggests that the biofilm– adhesive interaction exceeds the biofilm–substrate interaction more effectively than with ox-PDMS, thereby promoting the transfer of the biofilm from the glass substrate to the hydrophobic elastomer. The increasing dependence of removal efficiency on contact load indicates that a combination of interfacial and mechanical factors governs underwater biofilm peeling. Based on the significant difference between PDMS and ox-PDMS, adhesive peeling is dominated by interfacial interactions at contact loads ≤ 5 mN, whereas contact loads ≥ 10 mN generate sufficient mechanical stress to displace interfacial water and transfer or displace *S. aureus* biofilms from glass.^45,46^ Based on this initial observation, we next investigated whether underwater adhesives specifically designed to maximize interfacial contact and adhesion could further enhance peeling efficiency.

### 2.3 Hydrophobic adhesives require sufficient cohesive strength for mechanical stress transfer to enable adhesive peeling of *S. aureus* biofilms

Having established that hydrophobic interactions enable underwater biofilm peeling at low contact loads, we next investigated whether adhesives specifically designed for underwater adhesion could further enhance biofilm removal under these conditions. We synthesized HPE, a fluorescent, photocrosslinkable, hydrophobic polyester adhesive based on a previous report.^41,47^ The polyester contains two *N*-functional diol monomers: mOil provides hydrophobicity and low viscosity, and mPho enables UV-light-mediated crosslinking, allowing temporal control of the material properties of HPE (Figure 3A). We covalently tagged the HPE with rhodamine B to enable contact-interface imaging by confocal fluorescence microscopy during the adhesive peeling of biofilms.^48^

**Figure 3.**
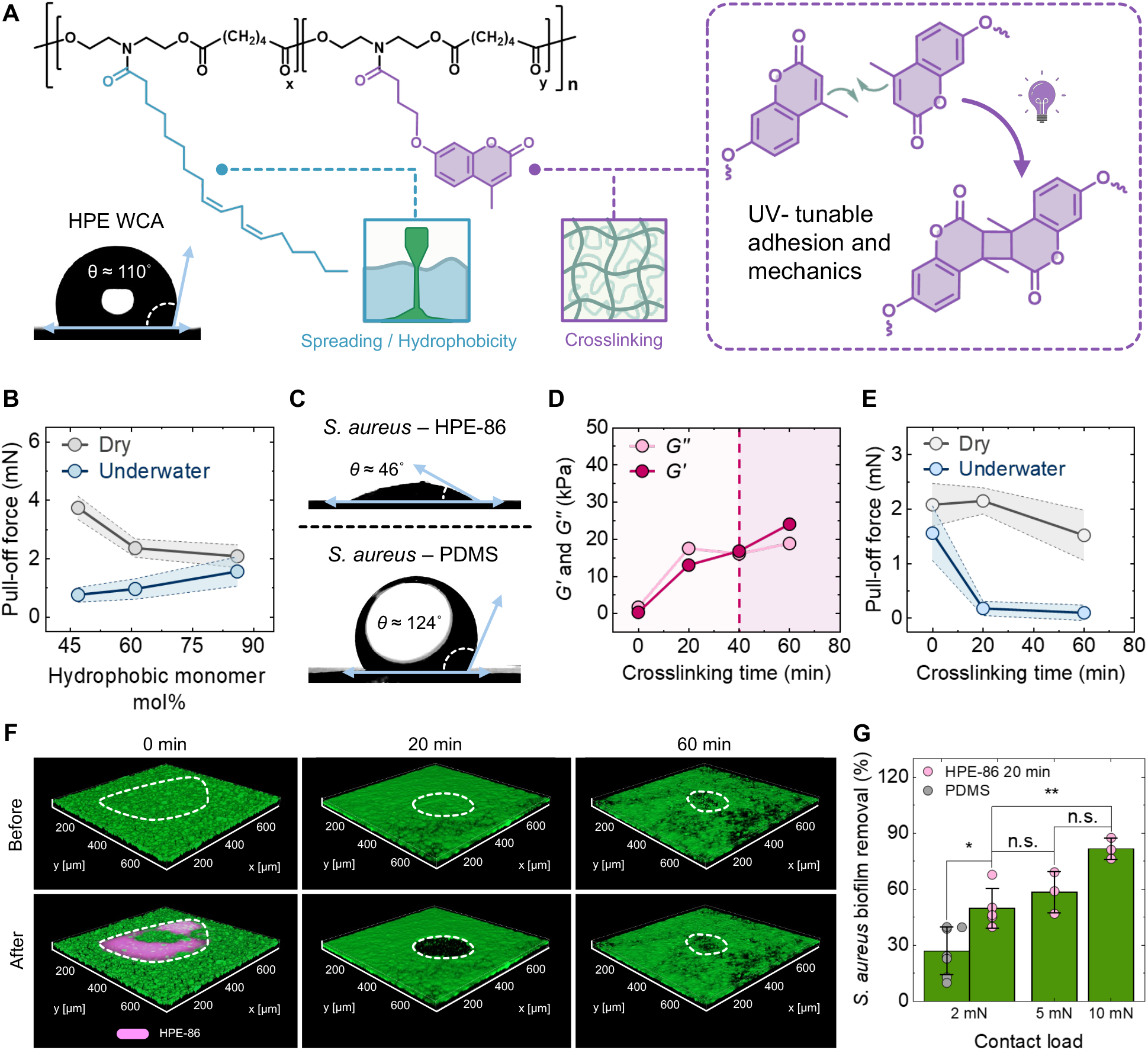
Hydrophobic polyester (HPE) underwater adhesive with tunable material properties used in this study to remove *S. aureus* biofilms at a lower contact load. (A) Chemical structure of HPE (WCA = 109.6 ± 1.8°) consisting of two monomeric units. The cyan monomer unit (mOil) contributes low viscosity, flow, and hydrophobicity to the polyester. The purple monomer unit (mPho) is a coumarin compound that provides tunable bulk moduli and adhesion strength for the polyester and is photocrosslinkable under UV light. (B) Composition-dependent pull-off force in dry and underwater conditions on glass. (C) Contact angles of the uncrosslinked HPE-86 and PDMS (Sylgard 184 Part A) on *S. aureus* biofilms, showing improved wetting behavior of HPE-86 compared with PDMS. (D) Storage (*G’*) and loss (*G’’*) moduli of HPE with 86 mol% of mOil (HPE-86) at strain = 0.1 % as a function of crosslinking time at 25 °C. (E) Dry and underwater pull-off force of HPE-86 at varying crosslinking times on glass. (F) Representative Z-stack images of *S. aureus* biofilm (green) and HPE-86 (pink) crosslinked for 0, 20, and 60 minutes (HPE-86 0 min, 20 min, 60 min) before and after peeling. The pink region at 0 min (after) indicates cohesive failure of HPE-86 0 min (Video S1). The highest biofilm removal was observed for HPE-86 20 min (Video S2), while HPE-86 60 min exhibited reduced removal due to adhesive failure at the biofilm–adhesive interface (Video S3). The HPE-86-coated PDMS lens was crosslinked externally before doing the adhesion tests and biofilm peeling. The biofilms were not exposed to UV light. (G) *S. aureus* biofilm removal for HPE-86 20 min at contact loads of 2, 5, and 10 mN. At 2 mN, HPE-86 20 min showed nearly two times higher *S. aureus* biofilm removal% compared with PDMS. Mean comparisons were performed using one-way ANOVA, and the significant differences across individual conditions were determined using Tukey’s test (n.s. = not significant, \**p* < 0.05, \*\**p* < 0.01, \*\*\**p* <0.001, \*\*\*\**p* <0.0001).

We first examined the effect that polymer composition (relative ratio between mOil and mPho) has on the bulk rheological and adhesive properties of HPE. As expected, increasing mOil content progressively reduced the zero-shear viscosity *η*_0_ of the polyester from 4.70 kPa⋅s (47 mol% of mOil) to 0.98 kPa⋅s (86 mol% of mOil) (Figure S11). The pull-off forces for each composition were subsequently evaluated under both dry and underwater conditions on glass substrates using a contact load of 2 mN and an equilibration time of 300 s (Figure 3B). Interestingly, the composition dependence of adhesion differed substantially between dry and underwater conditions. Under dry conditions, the HPE with the highest *η*_0_ composition exhibited the highest pull-off force at *F*_ad, dry_ = 3.74 ± 0.40 mN, whereas the lowest-viscosity composition exhibited the weakest pull-off force at *F*_ad, dry_ = 2.08 ± 0.39 mN (Figure 3B). In contrast, underwater adhesion followed the opposite trend, where the lowest-viscosity HPE demonstrated the highest pull-off force at *F*_ad, underwater_ = 1.56 ± 0.50 mN. Notably, the lowest-viscosity HPE showed similar pull-off force in both dry and underwater conditions, while higher-viscosity HPE adhesives showed a significant reduction in adhesion underwater. This observation suggests that hydrophobic polymers with higher chain mobility and conformability establish true molecular contact underwater in contrast to their higher-viscosity counterparts.^43^ Because the polyester with 86 mol% of mOil (HPE-86) exhibited the strongest underwater adhesion, it was selected for subsequent underwater biofilm peeling studies.

Based on the substantially higher underwater adhesion of HPE-86 compared with PDMS (PDMS-glass: *F*_ad,underwater_ ∼ 0 mN), we anticipated that HPE-86 would exhibit improved biofilm peeling efficiency. Towards this end, we first determined the theoretical thermodynamic work of adhesion (*w_ad_*) of the uncrosslinked PDMS (Sylgard 184 Part A) and HPE-86 using biofilm–adhesive contact angle experiments underwater. PDMS and HPE-86 showed contact angles of 123.8 ± 3.5° and 45.9 ± 7.9°, respectively (Figure 3C) with *S. aureus* biofilms. Using the Young-Dupré equation, we determined *w_ad_* = *γ_AW_* (1 + cos *θ_BA_*), where *γ_AW_* is the interfacial tension at the adhesive–water interface and *θ_BA_* is the underwater contact angle at the biofilm–adhesive interface. The *w_ad_* for PDMS and HPE-86 with *S. aureus* biofilms was 15.6 ± 2.3 mJ/m^2^ and 98.4 ± 5.7 mJ/m^2^, respectively, indicating stronger underwater interfacial interaction of HPE-86 with *S. aureus* biofilms compared with PDMS. However, during adhesive peeling of *S. aureus* biofilms, HPE-86 peeled only a negligible fraction ∼ 1.1 ± 2.7 % of the biofilm. The contact interface imaging (Video S1 and Figure 3F) revealed the presence of HPE-86 on the biofilm following peeling, indicating cohesive failure within the adhesive. In this regime, the adhesive established sufficient contact with the biofilm but lacked the cohesive strength required to transmit the mechanical stresses necessary for peeling. Consequently, failure occurred within the adhesive layer rather than at the biofilm–substrate interface or within the biofilm matrix.

The cohesive failure of HPE-86 during biofilm peeling therefore necessitated the crosslinking of the adhesive to increase its cohesive mechanical strength and enable the successful transfer of biofilms during peeling. HPE-86 was exposed to UV light (*λ* ∼ 365 nm, intensity ∼ 7 mW/cm^2^) and its mechanical properties were evaluated by performing strain amplitude sweep experiments (supporting information: section 4.2). At lower crosslinking time, the loss modulus (*G’’*) is higher than the storage modulus (*G’*) measured within the linear regime at 0.1 % strain, indicating that the material behaves as a viscous liquid (Figure 3D). As crosslinking time was increased, the *G’* gradually increased, and the material became increasingly solid-like. Beyond 40 minutes of crosslinking, *G’* exceeded *G’’*, indicating a transition to a predominantly solid-like state (Figure 3D).

The pull-off forces were determined for HPE-86 crosslinked for different time intervals. In dry conditions, the pull-off forces for HPE-86 0 min, 20 min, and 60 min were 2.08 ± 0.39, 2.15 ± 0.24, and 1.52 ± 0.46 mN, respectively (Figure 3E). The failure mode also changed with crosslinking, where uncrosslinked HPE-86 0 min failed cohesively, leaving residual polymer, while crosslinked HPE-86 failed adhesively, detaching cleanly from the glass. Adhesive failure from the biofilm-substrate interface is more desirable for peelable adhesives since no residual adhesive remains after peeling. In underwater conditions, HPE-86 0 min exhibited a measurable pull-off force of 1.56 ± 0.50, whereas the underwater pull-off forces for both HPE-86 20 min and HPE-86 60 min were negligible (Figure 3E). The failure modes remain similar to dry conditions, where HPE-86 0 min failed cohesively, while crosslinked HPE-86 (both 20 min and 60 min) failed from the glass-adhesive interface. The pull-force trends across dry and underwater conditions were notable since crosslinking did not significantly change adhesion strength in dry conditions, while underwater, crosslinking drastically weakened the adhesion strength, suggesting that the optimal degree of crosslinking for adhesion varies depending on the intervening medium. In dry conditions, the intervening medium (air) does not significantly interfere with contact between the adhesive and glass, whereas underwater, the adhesive with largely mobile chains (HPE-86 0 min) performs best, potentially because it can displace a significant amount of water from the interface, allowing adequate contact with the glass substrate underwater.

The adhesive peeling of *S. aureus* biofilms was tested using crosslinked HPE-86. For biofilm peeling at 2 mN contact load, HPE-86 20 min removed 49.8 ± 10.7 % of *S. aureus* biofilms, nearly twice the removal achieved by PDMS under similar conditions (Video S2 and Figure 3G). In contrast, HPE-86 60 min achieved only 6.0 ± 3.0 % removal (Video S3), indicating that excessive crosslinking limits the adhesive’s ability to establish true molecular contact with the biofilm. The greater removal by HPE-86 20 min likely results from an intermediate crosslinking density that provides sufficient cohesion for mechanical stress transfer, while still maintaining adequate conformability for true molecular contact with the biofilm. Further increasing the crosslinking time reduces biofilm peeling efficiency. As the adhesive becomes increasingly crosslinked, its ability to conform to the heterogeneous topography of the biofilm underwater is reduced, and effective contact formation becomes more challenging. At higher contact loads of 5 mN and 10 mN, HPE-86 20 min removed 58.3 ± 11.1 % and 81.6 ± 5.6 % of *S. aureus* biofilms, respectively (Figure 3G). These observations reveal three distinct peeling regimes: cohesive failure of the adhesive (HPE-86 0 min), successful biofilm transfer (HPE-86 20 min), and adhesive failure due to interfacial water-limited contact of the adhesive with biofilm (HPE-86 60 min). Thus, the successful underwater adhesive peeling of *S. aureus* biofilms emerges between these two failure extremes and requires a balancing act between interfacial adhesion and bulk cohesion.

### 2.4 Adhesive peeling of biofilms from hydrophobic surfaces is more challenging than from glass

The adhesive peeling framework indicates that biofilm removal is governed not only by biofilm– adhesive interactions but also by interactions between the biofilm and underlying substrates. Therefore, altering the substrate should influence the efficiency of peeling even when the adhesive and biofilm remain unchanged. To investigate the contribution of the biofilm–substrate interface, *S. aureus* biofilms were grown on substrates spanning a range of surface chemistries, including glass, trichloro(phenethyl)silane (PTS)-coated glass, and octadecyltrichlorosilane (OTS)-coated glass (Figure 4A). These substrates were selected to systematically vary surface functionalities while maintaining identical biofilm growth and peeling conditions. The adhesive peeling efficiency varied across the three substrates. For HPE-86 20 min, biofilm removal decreased as the substrate became increasingly hydrophobic (PTS and OTS), whereas PDMS and ox-PDMS exhibited no statistically significant differences in biofilm removal across the three substrates. For HPE-86 20 min, biofilm removal was 49.8 ± 10.7 %, 22.5 ± 6.5 %, and 21.4 ± 3.5 % for bare glass, PTS-coated glass, and OTS-coated glass, respectively (Figure 4B). This substrate-dependent reduction in biofilm removal is consistent with the well-established tendency of bacteria to preferentially adhere to hydrophobic surfaces over hydrophilic substrates such as glass, suggesting that stronger biofilm–substrate interactions on PTS-and OTS-coated glass contribute to reduced removal by HPE-86.^7,8,49–51^ The absence of a similar substrate-dependent response for PDMS and ox-PDMS suggests that their peeling performance is less sensitive to variations in biofilm– substrate interactions.

**Figure 4.**
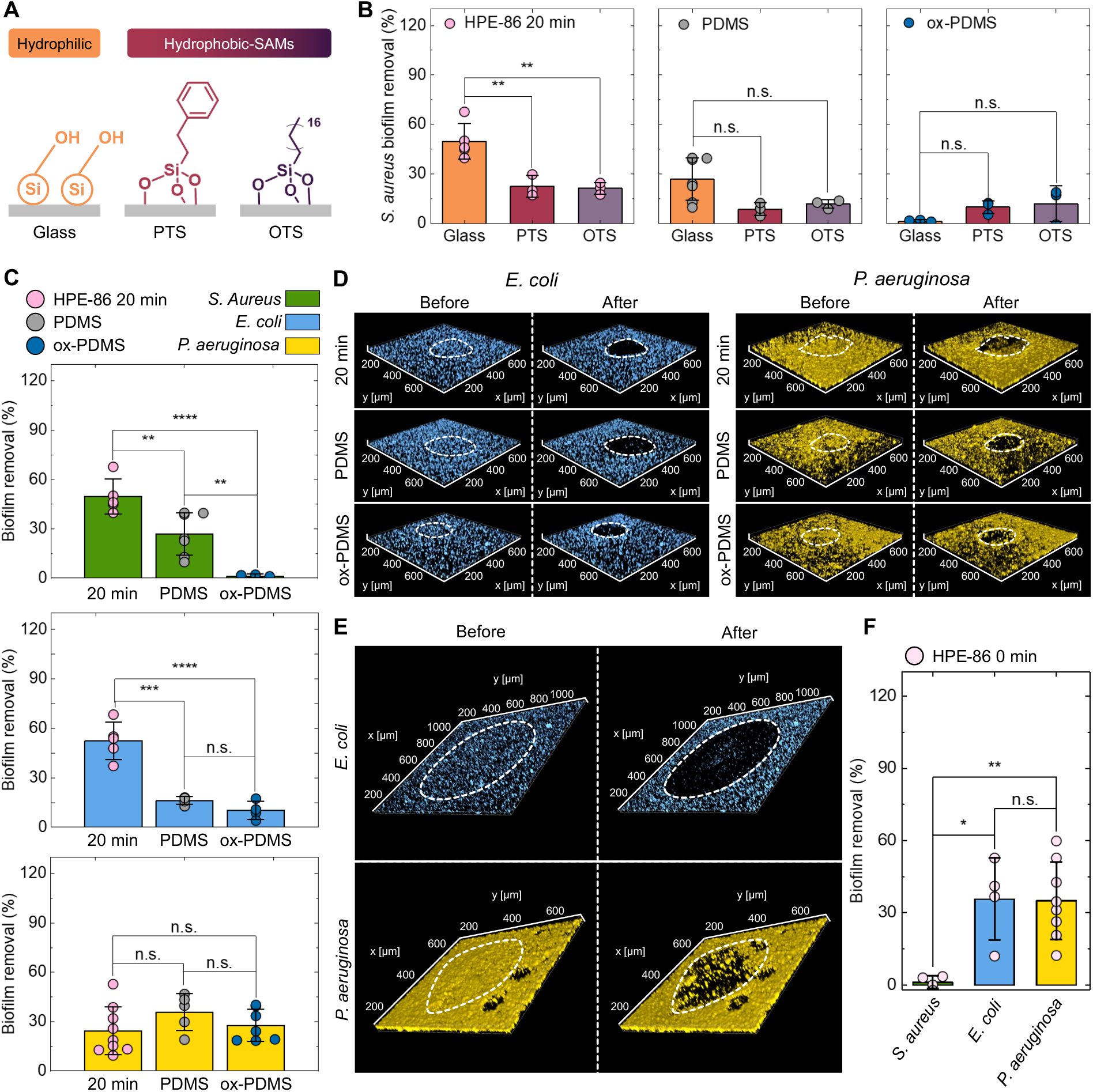
Influence of substrate and biofilm identity on adhesive peeling of biofilms. (A) Chemical structures of self-assembled monolayer (SAM)-coated surfaces with hydrophobic coatings: trichlorophenylsilane (PTS) and octadecyltrichlorosilane (OTS). (B) *S. aureus* biofilm removal by HPE-86 20 min, PDMS, and ox-PDMS on different surfaces. (C) Biofilm removal% of *S. aureus*, *Escherichia coli* (*E. coli*), and *Pseudomonas aeruginosa* (*P. aeruginosa*) by HPE-86 20 min (pink symbol), PDMS (gray symbol), and ox-PDMS (blue symbol) at a contact load of 2 mN. Bar height represents the mean, error bars show standard deviation, and the circular symbols show individual replicates. The HPE-86 20 min removed ∼ 50 % of the *E. coli* and *S. aureus* biofilms. All adhesives removed *P. aeruginosa* to a lower extent, at ∼ 30 %. (D) Representative 3D Z-stack images of *E. coli* and *P. aeruginosa* biofilms before and after peeling with each adhesive. Dashed white outlines indicate the contact region. (E) Representative images of adhesive peeling by HPE-86 0 min for *E. coli* (top, blue) and *P. aeruginosa* (bottom, yellow) biofilms. (F) HPE-86 0 min removed negligible *S. aureus* biofilm, while removing a significant percentage of *E. coli* and *P. aeruginosa* biofilms. Mean comparisons were performed using one-way ANOVA, and the significant differences across individual conditions were determined using Tukey’s test. (n.s. = not significant, \**p* < 0.05, \*\**p* < 0.01, \*\*\**p* <0.001, \*\*\*\**p* < 0.0001).

### 2.5 Biofilm species identity governs adhesive peeling efficiency and failure mechanism

Biofilms are molecularly and structurally diverse materials that differ in matrix composition, hydration state, architecture, interfacial adhesion, and mechanical properties. We hypothesized that biofilm peeling would depend not only on the adhesive and substrate, but also on the properties and identity of the biofilm itself. To examine the role of biofilm identity, we performed adhesive peeling experiments on biofilms formed by *E. coli*, *S. aureus*, and *Pseudomonas aeruginosa* (*P. aeruginosa*) using HPE-86 20 min, PDMS, and ox-PDMS. A low contact load of 2 mN was selected for these comparisons to minimize bulk mechanics and maximize sensitivity to differences in interfacial interactions.^42^ While substantially greater removal can be achieved at higher contact loads (Figure 3G), the low-load condition provides a more selective regime for comparing biofilm-dependent peeling.

HPE-86 20 min produced substantial removal across all three biofilms, with *E. coli* (52.5 ± 11.3 %) and *S. aureus* (49.8 ± 10.7 %) biofilms showing comparably high susceptibility to adhesive peeling, and *P. aeruginosa* generally exhibiting lower removal at 28.3 ± 18.3 % (Figure 4C and Figure 4D). PDMS and ox-PDMS produced distinct species-dependent removal profiles, and the rank order of peeling efficiency was not conserved across the different adhesives. Notably, HPE-86 0 min produced an interesting trend: it failed cohesively and showed negligible biofilm removal when interfaced with *S. aureus* biofilms, yet was able to remove *E. coli* (35.7 ± 17.1 %) and *P. aeruginosa* (35.1 ± 16.0 %) biofilms without cohesive failure (Figure 4E and Figure 4F). This species-dependent trend of peeling is consistent with the known diversity of biofilm extracellular matrices. Previous studies have shown that *S. aureus*, *E. coli*, and *P. aeruginosa* produce matrices that differ substantially in protein, polysaccharide, extracellular DNA, and water content.^52–58^ Such differences are expected to influence interfacial interactions with the adhesive as well as the mechanical response to the stress applied by the adhesive during peeling. Thus, biofilm identity does not simply scale the magnitude of removal but also alters the response to the contacting material. These results establish biofilm identity as an independent determinant of underwater adhesive peeling. The physicochemical origins of these differences, including the interfacial affinity and mechanical response of each biofilm, are examined quantitatively in the following section.

### 2.6 A four-body mechano-interfacial framework for adhesive peeling of biofilms underwater

The preceding experiments demonstrate that underwater biofilm peeling cannot be explained solely by a single adhesive property of the contacting material. The adhesive peeling of biofilms depended on adhesive surface chemistry, material properties of the adhesive, substrate surface chemistry, and biofilm identity. Collectively, these observations suggest that biofilm detachment emerges from a competition among four interacting bodies, namely, the adhesive (A: PDMS, ox-PDMS, or HPE), biofilm (B: *S. aureus*, *E. coli*, or *P. aeruginosa*), underlying substrate (S: glass, PTS, or OTS), and the intervening water phase (W: aqueous minimal media). The outcome of peeling is therefore governed by the interfacial affinity and mechanical stress associated with these living and non-living interfaces.

To formalize the interfacial component framework, we determined the biofilm–adhesive work of adhesion (*w_ad_*_,*BA*_) from the adhesive-biofilm contact angle using the Young-Dupré relation:

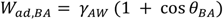

Where *γ_AW_* is the adhesive–water interfacial tension and *θ_BA_* is the contact angle between the biofilm and the uncrosslinked adhesive underwater. *w_ad_*_,*BA*_ explicitly incorporates the water phase because it describes the formation of the biofilm–adhesive interface from surfaces initially separated by water.

Similarly, a biofilm–adhesive mechanical mismatch parameter (*α_BA_*) was derived for describing the mechanical stress transfer associated with biofilm peeling,

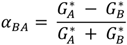

Where *G_A_*^∗^ and *G_B_*^∗^ are complex moduli of the adhesive and biofilm measured at the same strain amplitude. The details on the rheological measurements for biofilms can be found in the supporting information (section 5.4). When *α_BA_* → − 1, the adhesive is substantially softer than the biofilm, at *α_BA_* = 0, the adhesive and biofilm have matching *G*^∗^, and as *α_BA_* → + 1, the adhesive has low mechanical deformation compared with the biofilm. Note that due to experimental limitations, biofilm rheology was performed on isolated biofilms, and the measurements were recorded under humid conditions that are different from the submerged conditions used for biofilm peeling.

The mechano-interfacial map based on *w_ad_*_,*BA*_–*α_BA_* values of each combination of adhesives and biofilms showed three distinct peeling responses for *E. coli*, *S. aureus*, and *P. aeruginosa* (Figure 5A). For *E. coli*, biofilm removal was relatively insensitive to the mechanical properties of the adhesive polyester. HPE-86 0 min, 20 min, and 60 min have similar *w_ad_*_,*BA*_, but span a broad range of *α_BA_* from − 0.77 to 0.40 with *E. coli*. However, their biofilm removal efficiency remained consistent across crosslinking time (Figure 5B and Figure S21). Notably, HPE-86 0 min removed *E. coli* despite being considerably softer than the biofilm, indicating that once the *E. coli*–HPE interface is established, the adhesive does not require significant mechanical resistance for *E. coli* biofilm removal. For the *E. coli*–PDMS pair, PDMS combines a low *w_ad_*_,*BA*_ and a positive *α_BA_*, showing lower biofilm removal than the adhesive HPE-86 examples. These observations suggest that *E. coli* peeling is predominantly interfacially (*w_ad_*_,*BA*_) controlled. This behavior is consistent with the relatively weaker adhesion of planktonic *E. coli* on glass.^59–61^

**Figure 5.**
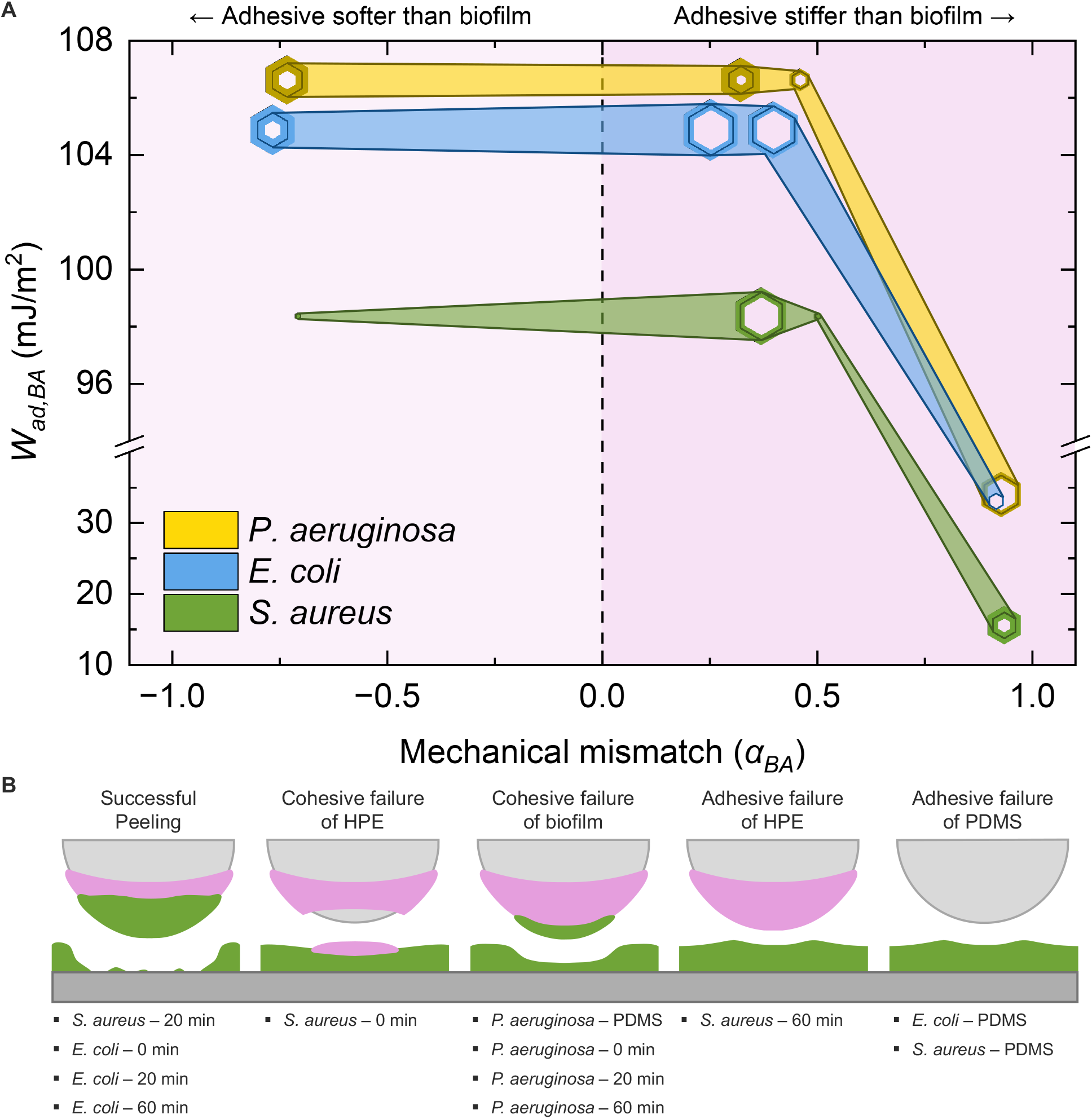
A mechano-interfacial map describing the biofilm removal trends observed during adhesive peeling. (A) Each hexagon symbol represents a biofilm–adhesive pair, which is plotted as a function of the work of adhesion between the biofilm and adhesive (*w_ad_*_,*BA*_), and the mechanical mismatch parameter (*α_BA_*). The background color distinguishes negative *α_BA_*, indicating that the adhesive is softer than the biofilm, from positive *α_BA_*, where the adhesive is stiffer than the biofilm. The diameter of each hexagon is proportional to the biofilm removal% of the particular biofilm-adhesive pair, with the interior hexagon outline marking the mean, and the halo shading around the outline corresponding to the standard deviation of the biofilm removal%. A wider shaded halo indicates higher standard deviation. The shaded band of the same color connects the data points among the same bacterial species peeled with different adhesives. The differences in the width of this shaded band when moving from left to right indicate how the mechanical properties and the interfacial interactions of the adhesive determine biofilm removal%. (B) Schematic of the peeling regimes observed for different biofilm–adhesive pairs. Successful peeling occurred when the adhesive bonded to the biofilm more strongly than the biofilm to the substrate. For example, this regime was observed for the HPE-86 20 min *S. aureus* peeling, suggesting that optimal cohesion was a requirement for removal. Cohesive failure of HPE-86 0 min occurred when the *w_ad_*_,*BA*_ was high, but the polyester lacked the necessary bulk cohesion and therefore remained on the biofilm after peeling. Cohesive failure of the biofilm was observed only for *P. aeruginosa*, with biofilm removal for all adhesives being comparably low.

The adhesive peeling of *S. aureus* exhibits a different response than *E. coli*. For *S. aureus*, neither high *w_ad_*_,*BA*_ nor *α_BA_* linearly influences the biofilm removal efficiency. Although HPE-86 0 min interacts strongly with *S. aureus* biofilms, its lower mechanical resistance results in limited removal and cohesive failure within the polyester (Figure 5B). At the other extreme, HPE-86 60 min and PDMS both exhibit low *S. aureus* biofilm removal efficiency (Figure 5B). HPE-86 20 min, with its high *w_ad_*_,*BA*_ and optimal *α_BA_*, provides an appropriate balance of interfacial interactions, mechanical resistance, and conformability, resulting in the highest biofilm removal. Therefore, maximum biofilm removal occurs at the intermediate region of the mechano-interfacial map. The requirement for a *w_ad_*_,*BA*_–*α_BA_* balance to achieve optimal *S. aureus* removal, contrasted with the effective removal of *E. coli* biofilms using the same, mechanically weak HPE-86 0 min, suggests that *S. aureus* biofilms exhibit stronger adhesion to glass than *E. coli* biofilms. Because we could not experimentally determine biofilm–glass adhesion with our experimental setup, the difference in response in the mechano-interfacial map can be interpreted as a relative difference in biofilm–substrate adhesion from the peeling response.

*P. aeruginosa* defines a third regime in the *w_ad_*_,*BA*_–*α_BA_* map. Across all tested adhesive conditions, spanning a broad range of *w_ad_* and *α_BA_*, biofilm removal remained comparably similar for *P. aeruginosa* biofilms (Figure 5B and Figure S21). For example, the mechanically weak, low-modulus HPE-86 0 min and the mechanically resistant, high-modulus PDMS achieved identical *P. aeruginosa* biofilm removal (Figure 5B). This adhesive-insensitive response in biofilm removal is consistent with a cohesively weak biofilm matrix. However, in the rheological measurement of biofilms, this mechanical fragility of *P. aeruginosa* was not evident when compared with *E. coli* or *S. aureus* (Figure S19). It is possible that the liquid-like behavior of *P. aeruginosa* is only revealed underwater. The adhesive peeling in this case likely represents the detachment of an upper portion of the biofilm while the basal layer remains attached to the substrate. This was also evident from the top-layer-biased removal of *P. aeruginosa* biofilm compared to *S. aureus* and *E. coli* (Figure S22). In this regime, the biofilm removal provides limited information about the strength of *P. aeruginosa* biofilm adhesion to glass because the biofilm cohesively fails before the underlying interface is challenged.

The mechano-interfacial map identifies three distinct behaviors in adhesive peeling of biofilms underwater: interfacially-controlled peeling for *E. coli*, coupled interfacial-mechanical peeling for *S. aureus*, and biofilm cohesion-limited removal for *P. aeruginosa*. Therefore, adhesive peeling does not simply report how strongly biofilm adheres, but where the weakest mechanical or interfacial pathway resides within the complete adhesive-biofilm-substrate-water four-body system.

## 3.0 DISCUSSION AND OUTLOOK

Inspired by the adhesive removal of biofilms observed in marine organisms, in this study, we developed an adhesive-enabled peeling of bacterial biofilms underwater. We found that hydrophobic adhesive polymers such as PDMS and HPE are fouling-release adhesives that can remove > 80 % biomass of bacterial biofilms underwater, corresponding to an areal treatment capacity of ∼ 100 cm^2^ per gram of adhesive. By varying the mechanical and interfacial properties of the adhesives and biofilms, we developed a mechano-interfacial map to study the trends of biofilm removal efficiency. An important consequence of this framework is that adhesive peeling provides a quantitative method for probing biofilm–substrate interactions by comparing differences in biomass loss patterns upon peeling. While current approaches for measuring biofilm adhesion remain challenging to implement underwater and are harsh on delicate biofilms, adhesive peeling in association with the *w_ad_*_,*BA*_–*α_BA_* map reports the outcome of the competition between biofilm adhesion and transfer. For example, the observation that identical biofilms exhibited markedly different removal efficiencies on surfaces with different chemistries indicates that biofilm removal% contains information regarding the strength of biofilm attachment to a given substrate. Adhesive peeling, therefore, provides a practical route for comparing biofilm retention across materials and evaluating antifouling or biofilm-resistant surfaces.

The species-dependent peeling behavior further demonstrates that adhesive peeling is sensitive to biofilm-specific physicochemical properties. Despite identical experimental conditions, *S. aureus*, *E. coli*, and *P. aeruginosa* exhibited distinct detachment responses. Because matrix composition, hydration state, and structural organization differ substantially among these species, adhesive peeling effectively converts these differences into a measurable transfer response. For example, between *E. coli, S. aureus*, and *P. aeruginosa*, the biofilm removal% response to polymer mechanics is notably different. The *w_ad_*_,*BA*_–*α_BA_* map distinguished these biofilms through their peeling behavior, suggesting that adhesive peeling may serve not only as a biofilm removal strategy but also as a functional assay for characterizing biofilm phenotypes and interfacial properties.

In summary, these results establish adhesive peeling as a quantitative framework for studying biofilm detachment underwater. By integrating adhesive interactions, hydration-mediated contact formation, substrate retention, and cohesive failure into a unified four-body mechano-interfacial map, adhesive peeling transforms biofilm removal from a technological challenge into a tractable interfacial mechanics problem. We anticipate that this framework will enable both the rational design of biofilm removal materials and a deeper understanding of attachment and detachment phenomena across complex submerged biological interfaces.

## 4.0 METHODS

### 4.1 Materials

All chemicals were used as received unless otherwise specified. Adipic acid (95 %) and rhodamine B (97 %) were purchased from Ambeed. Sylgard 184 was purchased from Fischer Scientific. LB broth and M9 minimal medium with 0.2 % glucose, were purchased from Teknova. *E. coli* (ATCC 25922) was purchased from American Type Culture Collection (ATCC). *S. aureus* (ATCC 25923) and *P. aeruginosa* (PAO 1) were kind gifts from Dr. Abraham Joy.

### 4.2 Synthesis of HPE

The polyesterification procedure was adapted from Gokhale et al.^62^ Two diol monomers, mOil and mPho (Supporting Information: sections 3.1 and 3.2), were reacted with adipic acid. Three hydrophobic polyesters (HPE) were prepared with varying mOil:mPho ratios. The synthesis of HPE-86 (target mOil:mPho ratio = 90:10) was carried out as follows. In a 100 mL round-bottom flask equipped with a magnetic stir bar, mOil (3.48 g, 9.46 mmol, 1.8 eq), mPho (0.3668 g, 1.05 mmol, 0.2 eq), adipic acid (1.54 g, 10.51 mmol, 2.0 eq), and 4-(dimethylamino)pyridinium 4-toluenesulfonate (1.24 g, 4.20 mmol, 0.8 eq) (DPTS) were added. The flask was then connected to a Schlenk line, dried under vacuum for 15 min, and backfilled with N_2_. The vacuum-N2 cycles were repeated three times. Under N_2_, 50 mL anhydrous DCM was added to the flask, and the mixture was cooled in an ice bath for 20 min. DIC (5.30 g, 42 mmol, 8.0 eq) was added dropwise, and the reaction was stirred for 48 h at room temperature. Upon completion, the mixture was filtered, and the filtrate was concentrated under reduced pressure and purified by precipitation in MeOH. The obtained HPE was dried under high vacuum overnight to obtain an isolated yield of ∼ 54 %. The targeted mOil:mPho compositions were 90:10, 60:40, and 50:50. The resulting compositions determined from ^1^H NMR were 86:14 (Figure S6), 61:39 (Figure S7), and 47:53 (Figure S8), respectively (HPE-86, HPE-61, and HPE-47). HPE-86 was fluorescently labeled when necessary, by adding rhodamine B (0.16 mg, 0.00031 mmol, 0.000063 eq) to the reaction mixture before DIC addition.

### 4.3 Preparation of hemispherical PDMS lenses and HPE coating

PDMS elastomers were prepared using Sylgard 184 (10:1 base-to-curing agent ratio) per the manufacturer’s protocol.^63^ The mixture was stirred for 5 min and degassed under vacuum for 30 min to remove air bubbles. After degassing, the mixture was preheated for 5 min at 80 °C and dispensed (∼ 50 µL) onto a glass substrate precoated with 1H,1H,2H,2H-perfluorodecyltrichlorosilane underwater. PDMS droplets were cured underwater at 70 °C for 24h. After curing, PDMS lenses (diameter ∼ 2.5 mm) were recovered and stored in a vacuum-sealed desiccator.

Prior to coating the HPE, PDMS lenses were treated with a corona torch for 30 s. A 4.3 wt% HPE solution in toluene was dip-coated onto the lenses for 30 s, keeping the lenses inverted to prevent flow-driven accumulation. Toluene was evaporated at room temperature for 2 hours, followed by drying under vacuum for 24 h. Approximately 600 μg of HPE is typically coated on the lenses. Samples were stored in a desiccator protected from light to avoid premature crosslinking.

### 4.4 Biofilm preparation

Biofilm culture was adapted from Vishwakarma et al.^48^ Bacterial cultures were first grown in LB medium under shaking at 37 °C for 24 h. Cultures were then centrifuged at 3000 rpm for 5 min, the supernatant was discarded, and the cell pellet was inoculated and resuspended in M9 minimal medium. The suspension was grown until reaching OD_600_ ∼ 0.5 and subsequently diluted in M9 medium to ∼10^6^ CFU mL^-1^. Approximately 800 µL of the diluted culture was added to each well of a 24-well glass-bottom plate or to a SAMs-coated glass coverslip placed inside a 24-well glass-bottom plate (supporting information: section 5.2). Plates were sealed with parafilm and incubated statically at 37 °C for 48 h. Following incubation, the supernatant was removed after centrifugation (3000 rpm, 5 min). To each well, 800 µL of new M9 medium was added. Biofilms were stained with DMAO for 30 min in the dark to a final DMAO concentration of 0.19 µM. Excess dye and planktonic cells were removed via pipette, and 0.85 % NaCl solution was added prior to imaging, per the manufacturer’s protocol.

### 4.5 Biofilm peeling experiments and imaging

The confocal microscopy imaging of the peeling process was recorded using 488 nm and 561 nm laser excitation. Before and after Z-stack images were acquired using identical settings (5 % laser power, 40 µm total thickness, 200 nm step size, exposure time). Different samples used varied exposure times. Background images containing only NaCl solution were acquired using the corresponding imaging settings and used for background subtraction during image quantification. The peeling experiments were performed using three probe types: (i) PDMS, (ii) surface-oxidized PDMS (ox-PDMS), and (iii) HPE-coated PDMS. ox-PDMS was prepared by treating PDMS with a corona torch for 30 s. For HPE-coated samples, PDMS was first corona-treated for 30 s to promote wetting, followed by dip-coating in the HPE solution (4.3 wt% in toluene) for 30 s. This coating was dried under ambient conditions for 2 h, followed by vacuum drying for 24 h to remove residual solvent. The probes were brought into contact with the biofilm at 1000 nm s^-1^ until the target contact load (2, 5, or 10 mN) was reached (Figure S17 and Figure 1B). This load was maintained for 5 min for equilibration, followed by probe retraction at 30 nm s^-1^.

### 4.6 Biofilm removal quantification

Biofilm removal was quantified using CellProfiler by comparing fluorescence intensity before and after peeling at the biofilm-adhesive contact area.^64^ The image analysis was restricted to the contact region segmented manually, defined as the area showing disruption in the DMAO-emission channel (PDMS or ox-PDMS) or rhodamine channel (HPE-86). Cohesive failure of the HPE-86 0 min with *S. aureus* was visualized using the rhodamine channel. All images used for biofilm removal quantification were analyzed in their native, unaltered state, without LUT adjustments or other intensity-modifying image processing.

### 4.7 Adhesion strength measurements

The PDMS lenses coated with HPE were brought into contact with glass at 1000 nm s^-1^ until the target contact load of ∼ 2 mN was reached. This load was maintained for 5 min for equilibration, followed by probe retraction at 30 nm s^-1^. Both dry and underwater measurements were recorded. In the underwater conditions, the substrate submerged in ultrapure water was brought in contact with the HPE-coated PDMS lenses. The pull-off force was determined from the peak force from the unloading curve (Figure S16).

### 4.8 Rheological characterization of PDMS, HPEs, and biofilms

Rheological measurements were performed on a TA Instruments Discovery HR-2 rheometer using a 20 mm parallel-plate geometry at 25 °C. For crosslinked HPE, the oscillatory strain amplitude sweep measurements were performed from 0.1 to 10 % strain at a constant angular frequency of 10 rad s^-1^ (Figure S13). Across all crosslinking times, *G’* and *G’’* remained constant within the linear viscoelastic region up to ∼ 2 % strain. For PDMS, the oscillatory strain sweep measurements were performed from 0.1 to 100 % strain and showed predominantly elastic solid behavior (Figure S14).

The biofilm samples used for rheological measurements were aseptically inoculated onto tryptic soy agar plates and incubated for 48 h at 37 °C. The visible biofilm colonies (diameter ∼ 0.50 to 1.00 mm) were individually collected and transferred to a 24-well plate.^22^ Plates were centrifuged at 1000 rpm for 5 min to distribute the biofilm evenly in the wells before rheological testing. An 8 mm parallel plate geometry at 25 °C (Figure S18) was used for performing the strain amplitude sweeps at a constant angular frequency of 1 rad s^-1^ over a range of 0.01 to 100 % strain (Figure S19). At least three replicates were performed for this analysis.

### 4.9 Statistical analysis

OriginPro was used for statistical analysis. Raw, unindexed input data were used for all statistical comparisons. Differences among groups were evaluated using one-way ANOVA, followed by Tukey’s post hoc test for pairwise mean comparisons. Homogeneity of variances was assessed using Levene’s test. Statistical significance was set at *p* < 0.05.

## 5.0 ACKNOWLEDGEMENTS

This work was supported by A.N.’s start-up funding from the University of Florida. The University of Florida’s Polymer Chemistry Laboratory and the Center for Nuclear Magnetic Resonance Spectroscopy in the Department of Chemistry are acknowledged for instrumentation. University of Florida Marston Library Makerspace 3D printing is acknowledged. Authors thank Evangelos Gatzogiannis for discussions regarding adhesive peeling instrumentation. Authors thank Profs. Austin M. Evans, Valeria D. Kleiman, and Brent S. Sumerlin for instrumentation.

During the preparation of this manuscript, chatgpt.com and claude.ai were used to enhance the readability and clarity of certain sections of the text. All content produced with the assistance of these tools was carefully reviewed, verified, and edited by the authors, who take full responsibility for the final version of the publication.

## 6.0 AUTHOR CONTRIBUTIONS

A. L. K. O and A. J. B. contributed equally. A. N. conceptualized this project. A. L. K. O. and A. J. B. designed and carried out the experiments. L. A. C. V., S. K. D., and Muskan helped with the polymer characterization. A. N. supervised the project. The manuscript was written by A. L. K. O., A. J. B., and A. N., with edits and input from L. A. C. V., S. K. D., and Muskan.

## Supporting Information

## 1 : Materials

All chemicals were used as received unless otherwise specified. *p*-Toluenesulfonic acid monohydrate (PTSA, 99 %), 4-(dimethylamino)pyridine (DMAP, 99 %), methyl 4-bromobutyrate (95 %), and *N*,*N*′-diisopropylcarbodiimide (DIC, >99 %) were purchased from Oakwood Chemical. 18-crown-6 (98 %), adipic acid (95 %), trichloro(phenethyl)silane (PTS, 95 %), and rhodamine B (97 %) were purchased from Ambeed. Diethanolamine (DEA, >99 %) was purchased from Chem-Impex. Sodium methoxide (NaOMe, >99 %), soybean oil (SBO), anhydrous potassium carbonate (K_2_CO_3_, 99 %), the Sylgard 184 silicone elastomer kit, and octadecyltrichlorosilane (OTS, 95 %) were purchased from Fisher Scientific.

Chloroform-*d* (CDCl_3_, 99.9 %) was purchased from Cambridge Isotope Laboratories, Inc. Analytical-grade organic solvents, including dichloromethane (DCM), *N*,*N*-dimethylformamide (DMF), ethyl acetate (EtOAc), methanol (MeOH), tetrahydrofuran (THF), dimethyl sulfoxide (DMSO), ethanol (EtOH), and toluene were obtained from Fisher Scientific.

*E. coli* (ATCC 25922) was purchased from American Type Culture Collection (ATCC). *S. aureus* (ATCC 25923) and *P. aeruginosa* (PAO 1) were kind gifts from Dr. Abraham Joy.^1^ Sterile media, including LB broth and M9 minimal medium with 0.2 % glucose, were purchased from Teknova. Sterile tryptic soy agar plates, used in the biofilm rheology study, were purchased from Teknova. DMAO (2 mM solution in DMSO) was purchased from Biotium. 24-well glass bottom plates were purchased from Cellvis, and glass coverslips (14 mm diameter, 0.16-0.19 mm thickness) were purchased from Chemglass Life Sciences. Stainless steel 90-degree angle needles, used for captive bubble experiments, were purchased from McMaster-Carr. Skybasic’s wireless digital microscope camera with 50–1000 × magnification was purchased from Amazon and used for contact angle measurements.

## 2 : Instrumentation

### 2.1 Nuclear magnetic resonance (NMR) spectroscopy

^1^H NMR spectra were acquired at 25 °C on a Bruker 400 MHz spectrometer using deuterated chloroform (CDCl_3_) as the solvent. Chemical shifts (*δ*) are reported in parts per million (ppm) and were referenced to the residual solvent signal of CDCl_3_ (*δ* = 7.26 ppm).

### 2.2 Size exclusion chromatography (SEC)

SEC was performed using an Agilent 1260 Infinity series liquid chromatography system equipped with an isocratic pump, autosampler, column compartment, and differential refractive index detector. Separation was achieved using two Waters Styragel HR-4E GPC columns (7.8 × 300 mm, 5 µm) in series with a preceding guard column (4.6 × 30 mm, 20 µm). Measurements were conducted at 40 °C using HPLC-grade THF as the mobile phase at a flow rate of 1.0 mL min^-1^ and an injection volume of 100 μL. The system was calibrated using narrow-dispersity polystyrene standards. Samples were prepared at 2 mg mL^-1^ and allowed to dissolve for 24 h prior to filtration through a 0.22 μm syringe filter.

### 2.3 Differential scanning calorimetry (DSC)

Thermal analysis was performed using a TA Instruments Discovery 2500 DSC with an RSC 90 refrigerated cooling system. Samples (5.0 ± 0.5 mg) were sealed in hermetic aluminum pans and analyzed against an empty reference pan. Under a nitrogen atmosphere (50 mL min^-1^), samples underwent two cooling-heating-cooling cycles from − 75 to 120 °C at a rate of 5 °C min^-1^ with 5 min isothermal holds at each temperature extreme. Data acquisition and analysis were performed using TRIOS software (v5.1.1).

### 2.4 Rheology

Rheological analysis was performed using a TA Instruments Discovery HR-2 rheometer with 8 mm or 20 mm parallel plate geometries. Data acquisition and analysis were performed using TRIOS software (v5.1.1).

### 2.5 Custom-built force testing system

The adhesion measurements and adhesive peeling were performed using a custom-built apparatus developed in our laboratory. This system is equipped with FUTEK’s 50 g capacity LSB200 load cell, with vertical (Z-axis) motion driven by a Thorlabs PIA13 piezo inertia actuator controlled by a KIM001 piezo inertia motor controller. The adhesion probe consisted of a knurled-head thumb screw (M3.5 × 0.6 mm thread size, 31 mm length) from McMaster-Carr attached to the active end of the load cell.

For each measurement, the probe was advanced toward the sample at 1000 nm s^-1^ until the target contact load (2, 5, or 10 mN) was reached. This load was maintained for 300 s to allow equilibration. The probe was then retracted at 30 nm s^-1^. During biofilm peeling experiments, the system was attached to a confocal microscope, enabling real-time visualization of the biofilm– adhesive contact, equilibration, and peeling as shown in Figure S1. Thorlabs Kinesis (v. 1.14.55) software was used to control the motor, while the force-time curve was monitored and acquired using FUTEK’s SENSIT software.

**Figure S1.**
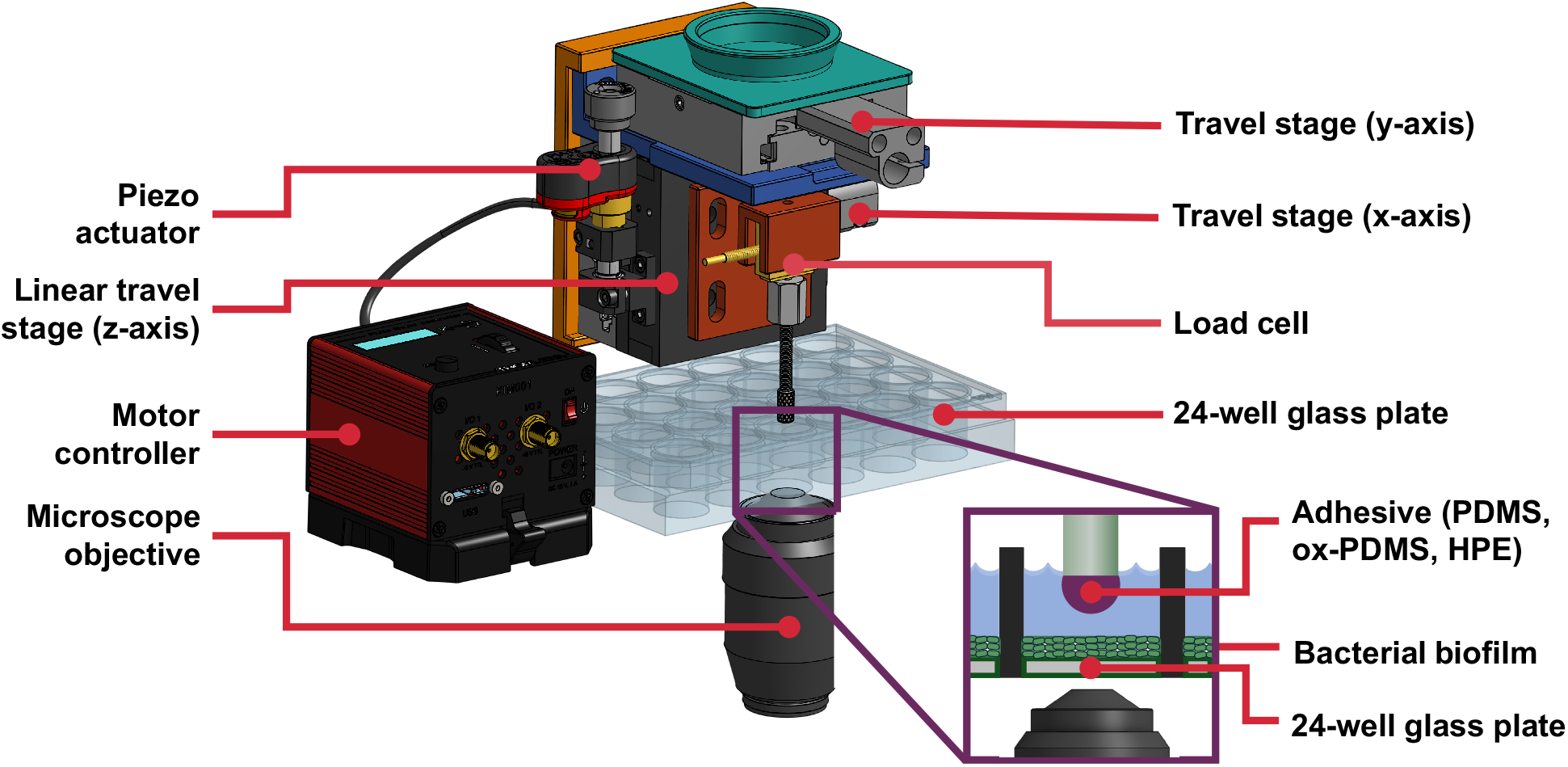
Custom-built force measurement coupled to a confocal microscope.

### 2.6 Spinning disk confocal microscopy

Fluorescence confocal microscopy imaging was performed using a Crest X-Light V3 spinning disk on a Nikon Ti2-Eclipse inverted microscope with Celesta light engine solid-state lasers. DMAO (biofilm stain) and Rhodamine B (polymer label) were excited using 488 nm and 561 nm lasers, respectively. A 10 × air objective (NA = 0.45) was used for imaging. All images were acquired using identical acquisition settings. Images were captured using a Kinetix sCMOS 10 MP camera. The Z-stacks were collected over a ± 20 µm range from the focal plane with a 200 nm step size. The image acquisition and processing were performed using NIS-Elements software. The background-corrected raw images were used for image analysis and the images and videos shown in the Figures are processed using denoise.ai in NIS-Elements software.

## 3 : Synthetic Methods

### 3.1 Synthesis of hydrophobic diol monomer (mOil)

The hydrophobic diol monomer was synthesized using a similar procedure as reported previously with minor modifications.^2^ The synthetic route is shown in Scheme S1. DEA (16.40 g, 156 mmol, 3.0 eq), SBO (43.40 g, 52 mmol, 1.0 eq), and NaOMe (0.281 g, 5.2 mmol, 0.1 eq) were combined in a 500 mL round-bottom flask. The mixture was heated and stirred at 90 °C under vacuum for 6 h, with reaction progress monitored by thin-layer chromatography (TLC). Upon completion, the crude product was purified by column chromatography using 90:10 (DCM:MeOH) as the eluent. The purified product was dried under reduced pressure overnight to remove residual solvent (Figure S2).

**Scheme S1.**
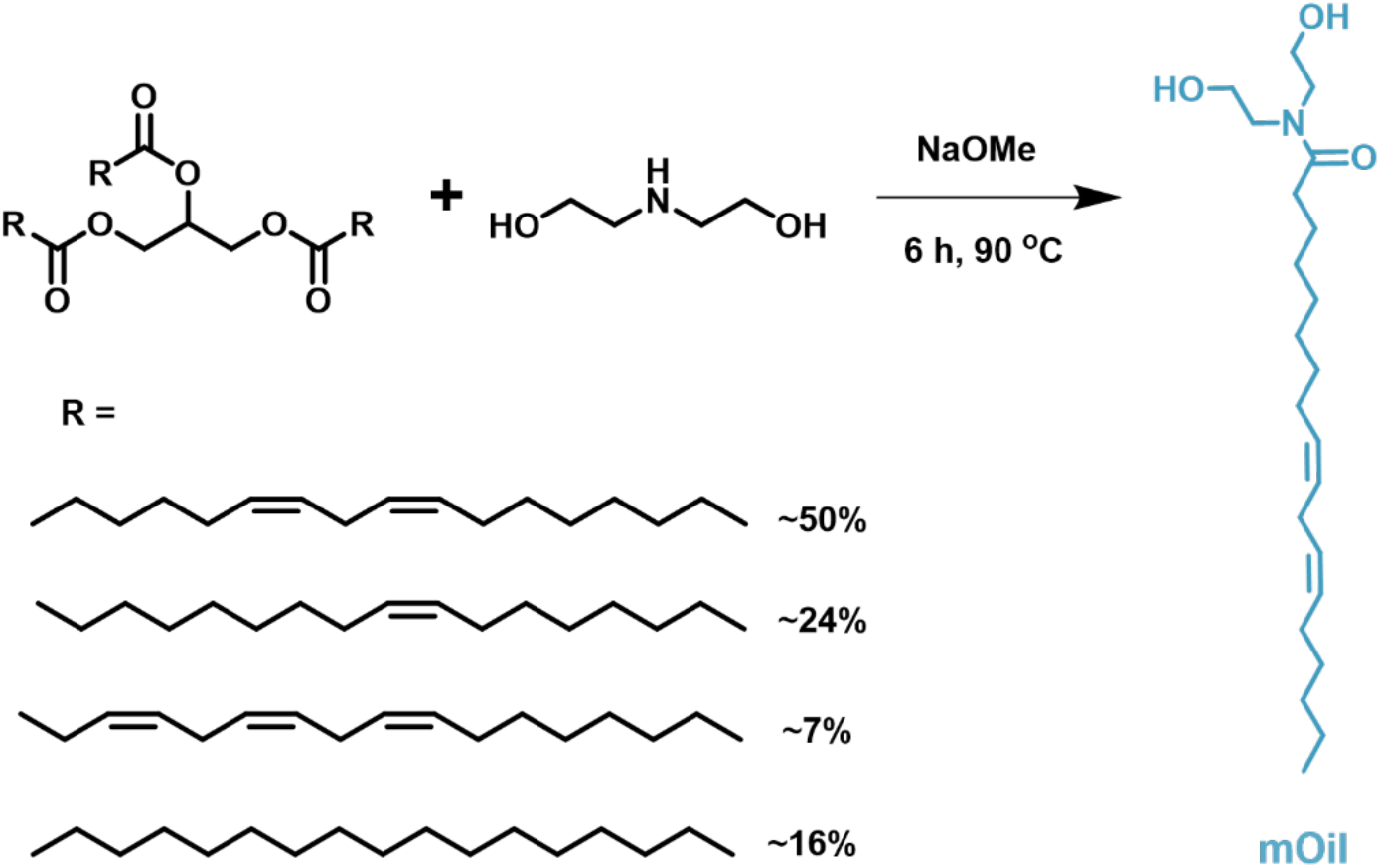
Schematic representation of the synthesis of the hydrophobic diol monomer (mOil).

**Figure S2.**
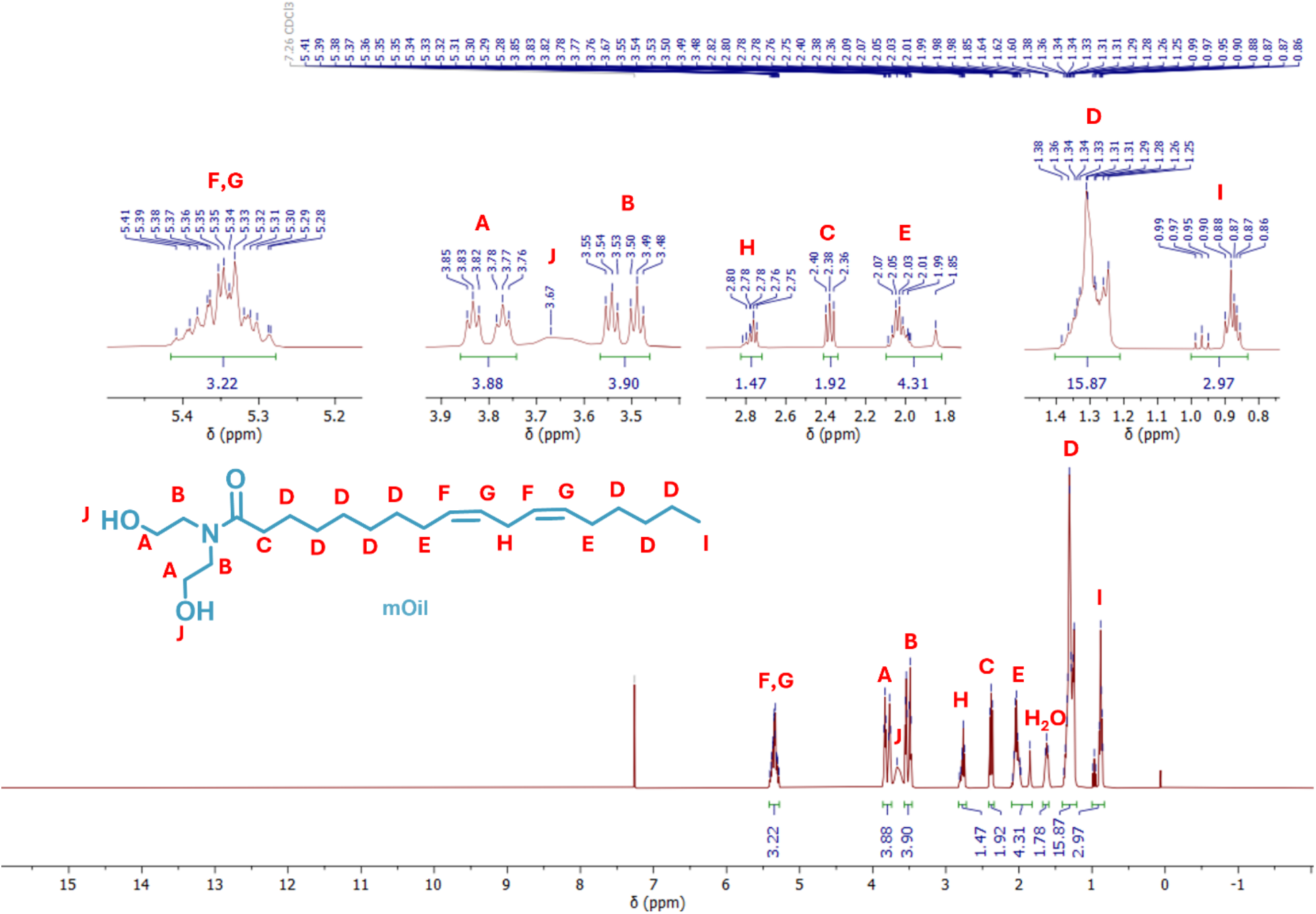
^1^H NMR spectrum of the hydrophobic diol monomer in CDCl_3_.

### 3.2 Synthesis of photocrosslinkable diol monomer (mPho)

The synthetic route involves a two-step reaction shown in Scheme S2 and was adapted from the procedure reported by Govindarajan et al.^3^

#### Step 1. Synthesis of methyl 4-((4-methyl-2-oxo-2H-chromen-7-yl)oxy)butanoate

4-Methylumbelliferone (15.94 g, 91 mmol, 1.0 eq), anhydrous K_2_CO_3_ (25.04 g, 181 mmol, 2.0 eq), 18-crown-6 (1.22 g, 4.61 mmol, 0.05 eq), and methyl 4-bromobutyrate (22.9 mL, 181 mmol, 2.0 eq) were combined in a 500 mL round-bottom flask. DMF (∼ 250 mL) was added to dissolve the reagents, and the reaction mixture was stirred at 80 °C overnight while protected from light with aluminum foil. Upon completion, the reaction mixture was filtered, and the solvent was removed under reduced pressure. Residual DMF was removed by liquid-liquid extraction with water and EtOAc. The organic layer was dried over Na_2_SO_4_, concentrated under reduced pressure, and purified by column chromatography using 90:10 (DCM:MeOH) as the eluent. The product was dried under high vacuum overnight to afford the desired compound in 59 % yield (Figure S3).

**Scheme S2.**
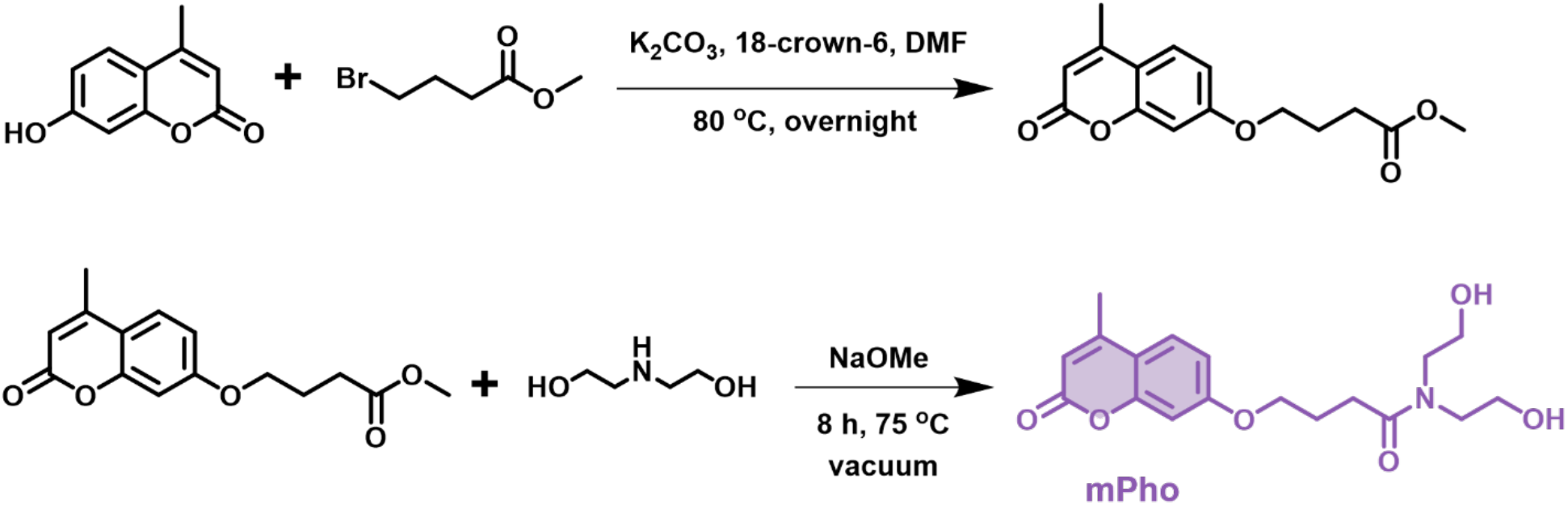
Schematic representation of synthesis of photocrosslinkable diol monomer (mPho).

**Figure S3.**
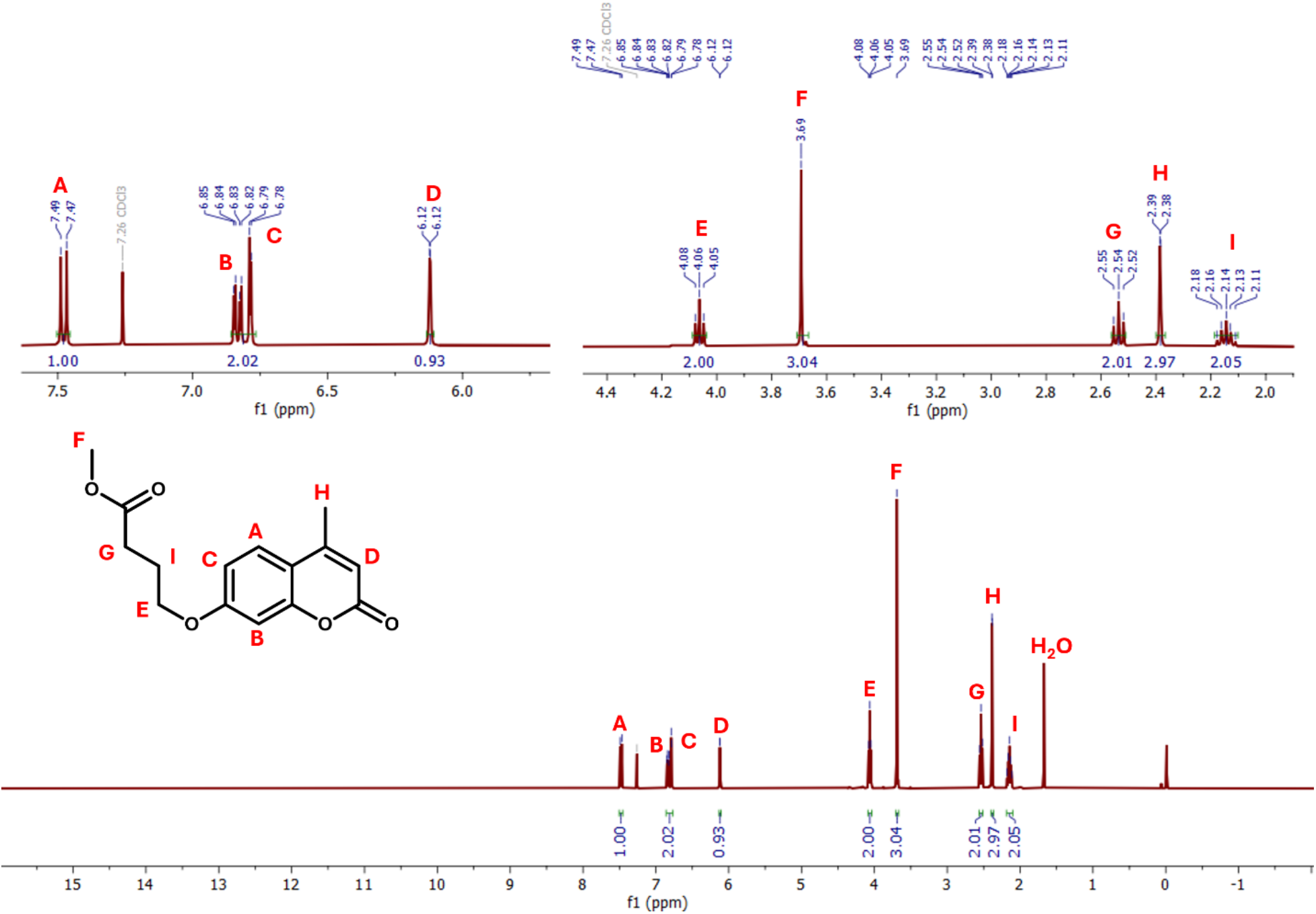
^1^H NMR spectrum of methyl 4-((4-methyl-2-oxo-2H-chromen-7-yl)oxy)butanoate in CDCl_3_.

#### Step 2. Synthesis of N,N-bis(2-hydroxyethyl)-4-((4-methyl-2-oxo-2H-chromen-7 yl)oxy)butanamide

Methyl 4-((4-methyl-2-oxo-2H-chromen-7-yl)oxy)butanoate (27.12 g, 98.27 mmol, 1.0 eq), DEA (42.36 g, 402.9 mmol, 4.1 eq), and NaOMe (0.265 g, 4.91 mmol, 0.05 eq) were combined in a round-bottom flask and stirred at 75 °C under vacuum for 8 h. The crude product was purified by column chromatography using 90:10 (DCM:MeOH) as the eluent and dried under high vacuum overnight to yield the final product in 90 % yield (Figure S4).

^1^H NMR (400 MHz, CDCl_3_) *δ* 7.47 (d, *J* = 8.8 Hz, 1H), 6.86 – 6.78 (m, 2H), 6.11 (d, *J* = 1.2 Hz, 1H), 4.08 (t, *J* = 6.1 Hz, 2H), 3.86 (t, *J* = 5.0 Hz, 2H), 3.79 (t, *J* = 5.2 Hz, 2H), 3.58 – 3.52 (m, 4H), 2.64 (t, *J* = 7.1 Hz, 2H), 2.38 (d, *J* = 1.3 Hz, 3H), 2.17 (p, *J* = 6.6 Hz, 2H).

**Figure S4.**
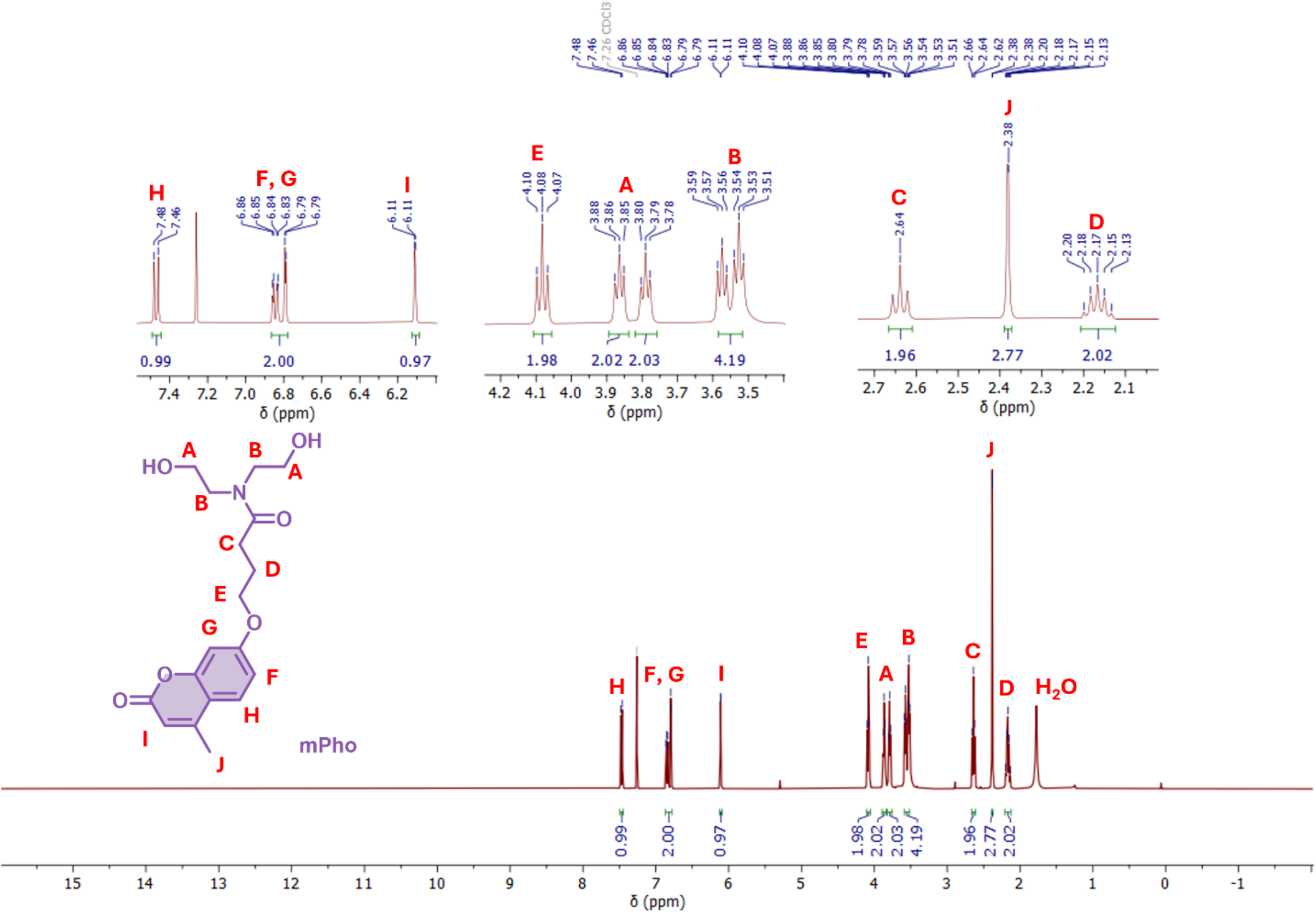
^1^H NMR spectrum of N,N-bis(2-hydroxyethyl)-4-((4-methyl-2-oxo-2H-chromen-7-yl)oxy)butanamide in CDCl_3_.

### 3.3 Synthesis of 4-(dimethylamino)pyridinium 4-toluenesulfonate (DPTS) catalyst

DMAP (20.00 g, 163.7 mmol, 1.0 eq) and *p*-toluenesulfonic acid monohydrate (31.14 g, 163.7 mmol, 1.0 eq) were dissolved in THF (∼ 250 mL) in a 500 mL round-bottom flask.^4^ The reaction was stirred at room temperature, during which the product precipitated. The solid was collected by filtration and dried under high vacuum overnight to afford the DPTS catalyst in 97 % yield (Figure S5).

^1^H NMR (400 MHz, CDCl_3_) *δ* 8.26 (d, *J* = 7.6 Hz, 2H), 7.85 (d, *J* = 8.3 Hz, 2H), 7.18 (d, *J* = 7.9 Hz, 2H), 6.72 (d, *J* = 7.7 Hz, 2H), 3.23 (s, 6H), 2.35 (s, 3H).

**Figure S5.**
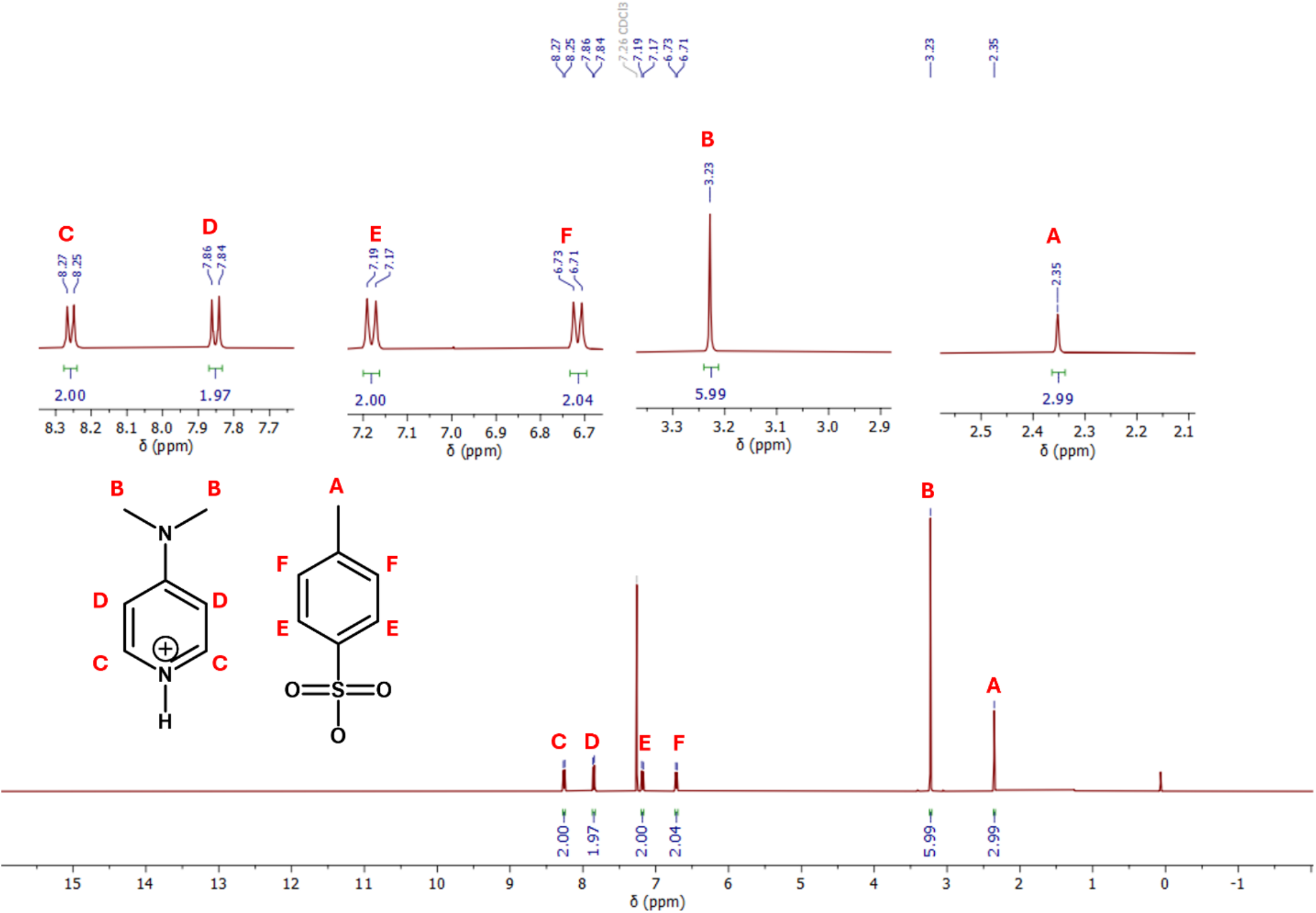
^1^H NMR spectrum of DPTS catalyst in CDCl_3_.

### 3.4 Synthesis of hydrophobic polyesters (HPE)

The polyesterification procedure as shown in Scheme S3 was adapted from Gokhale et al.^5^ The reaction employed the diol monomers mOil and mPho (synthesized above) and adipic acid as the dicarboxylic acid monomer. Three hydrophobic polyesters (HPE) were prepared with varying mOil:mPho ratios.

As an example, a target 90:10 mOil:mPho composition was prepared using mOil (3.48 g, 9.46 mmol, 1.8 eq), mPho (0.3668 g, 1.05 mmol, 0.2 eq), adipic acid (1.54 g, 10.51 mmol, 2.0 eq), and DPTS (1.24 g, 4.20 mmol, 0.8 eq, 8 eq) in a 100 mL round-bottom flask equipped with a magnetic stir bar. For fluorescence labeling in biofilm peeling experiments, rhodamine B was added at 0.000063 eq (0.16 mg, 0.00031 mmol).

The flask was connected to a Schlenk line, dried under vacuum for 15 min, and backfilled with N_2_ (three cycles). Anhydrous DCM (50 mL) was added under nitrogen, and the mixture was cooled in an ice bath for 20 min. DIC (5.30 g, 42 mmol, 8.0 eq) was then added dropwise, and the reaction was stirred for 48 h. Upon completion, the mixture was filtered, and the filtrate was concentrated and precipitated in MeOH. The HPE was reprecipitated four times to remove residual DIC and byproducts, then dried under high vacuum overnight. The isolated yield was ∼ 54 %. The targeted mOil:mPho compositions were 90:10, 60:40, and 50:50. The resulting compositions determined from ^1^H NMR were 86:14 (Figure S6), 61:39 (Figure S7), and 47:53 (Figure S8), respectively (HPE-86, HPE-61, and HPE-47).

**Scheme S3.**
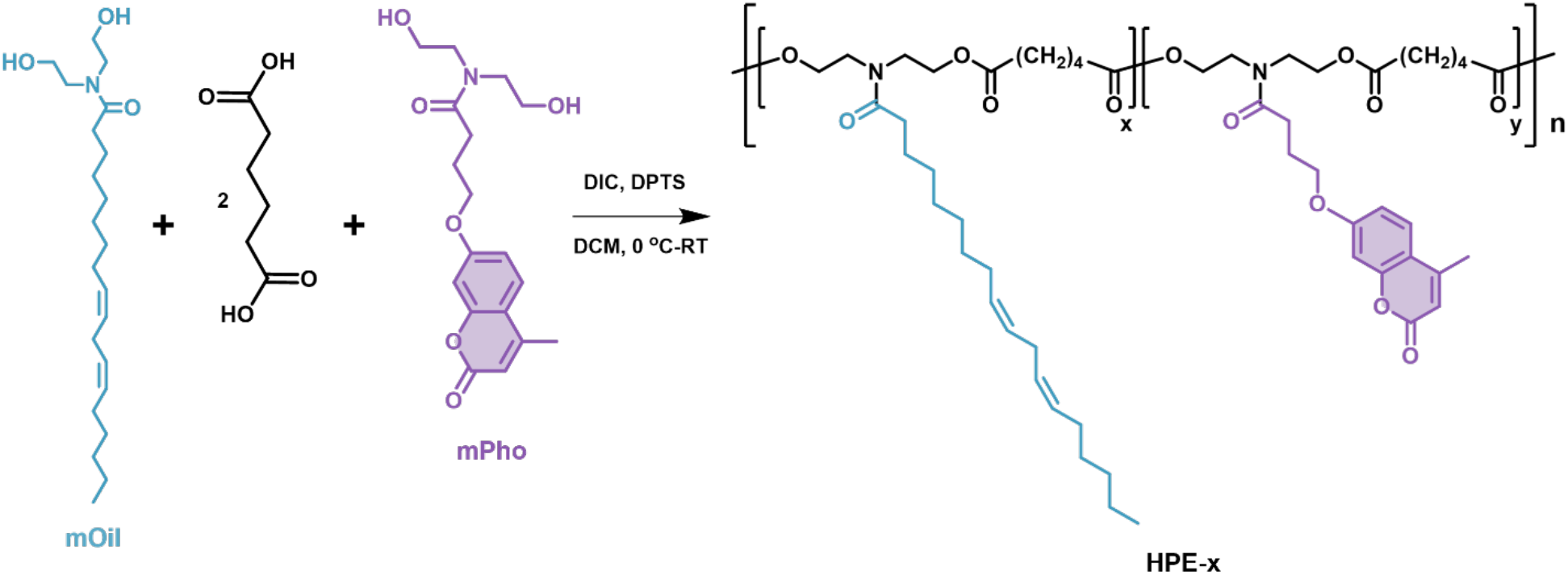
Schematic representation of polyesterification.

**Figure S6.**
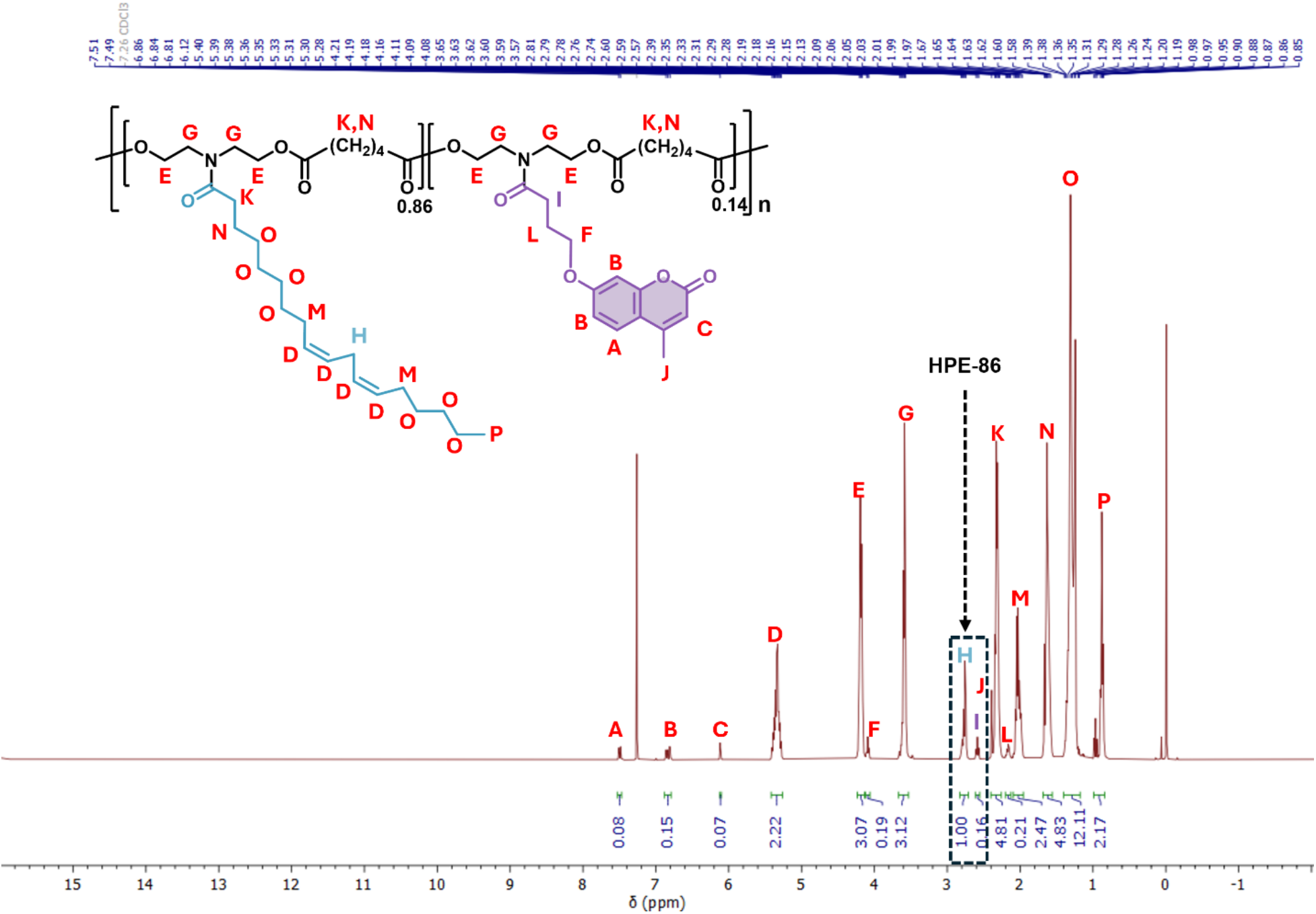
^1^H NMR spectrum of HPE-86 in CDCl_3_.

**Figure S7.**
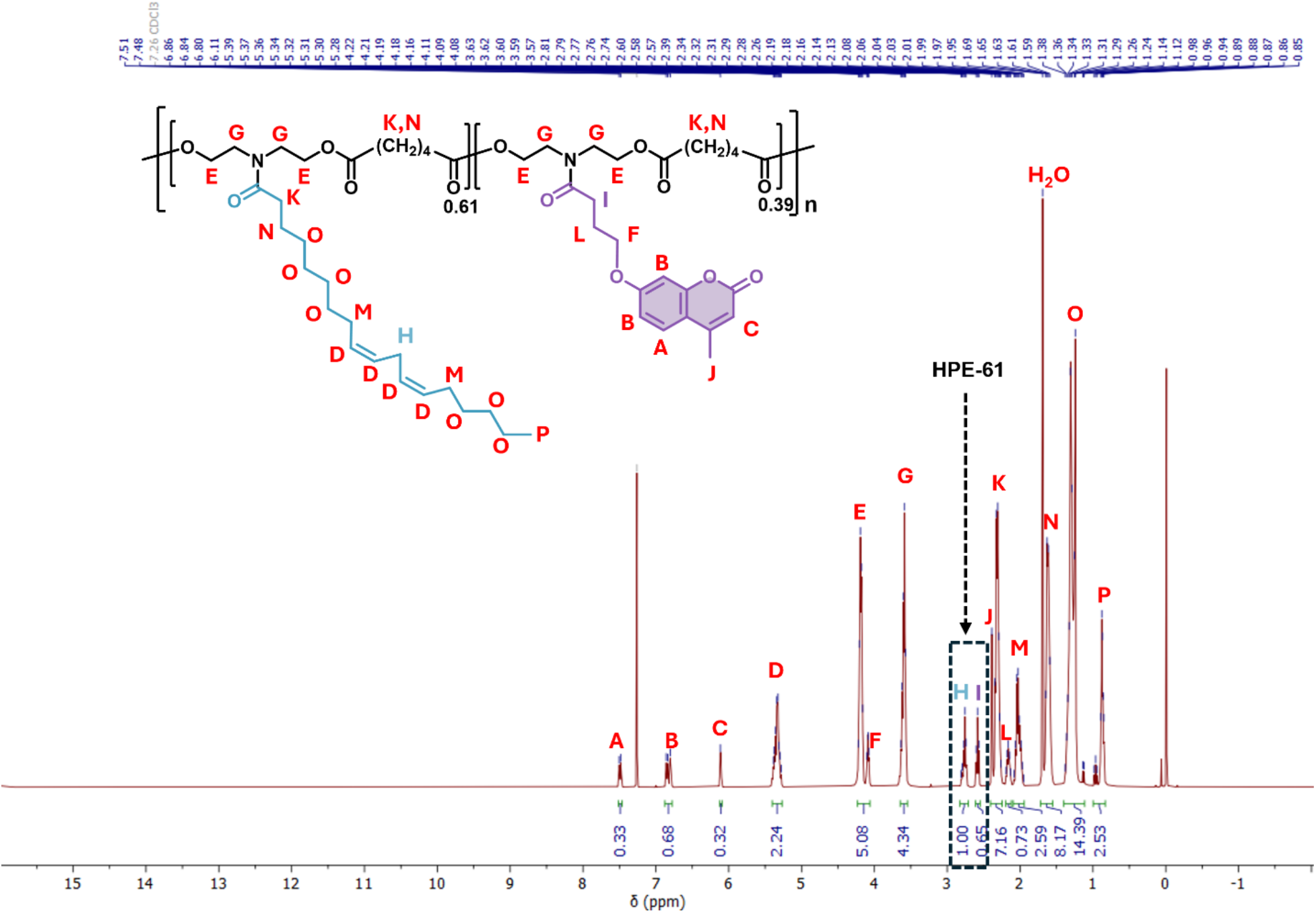
^1^H NMR spectrum of HPE-61 in CDCl_3_.

**Figure S8.**
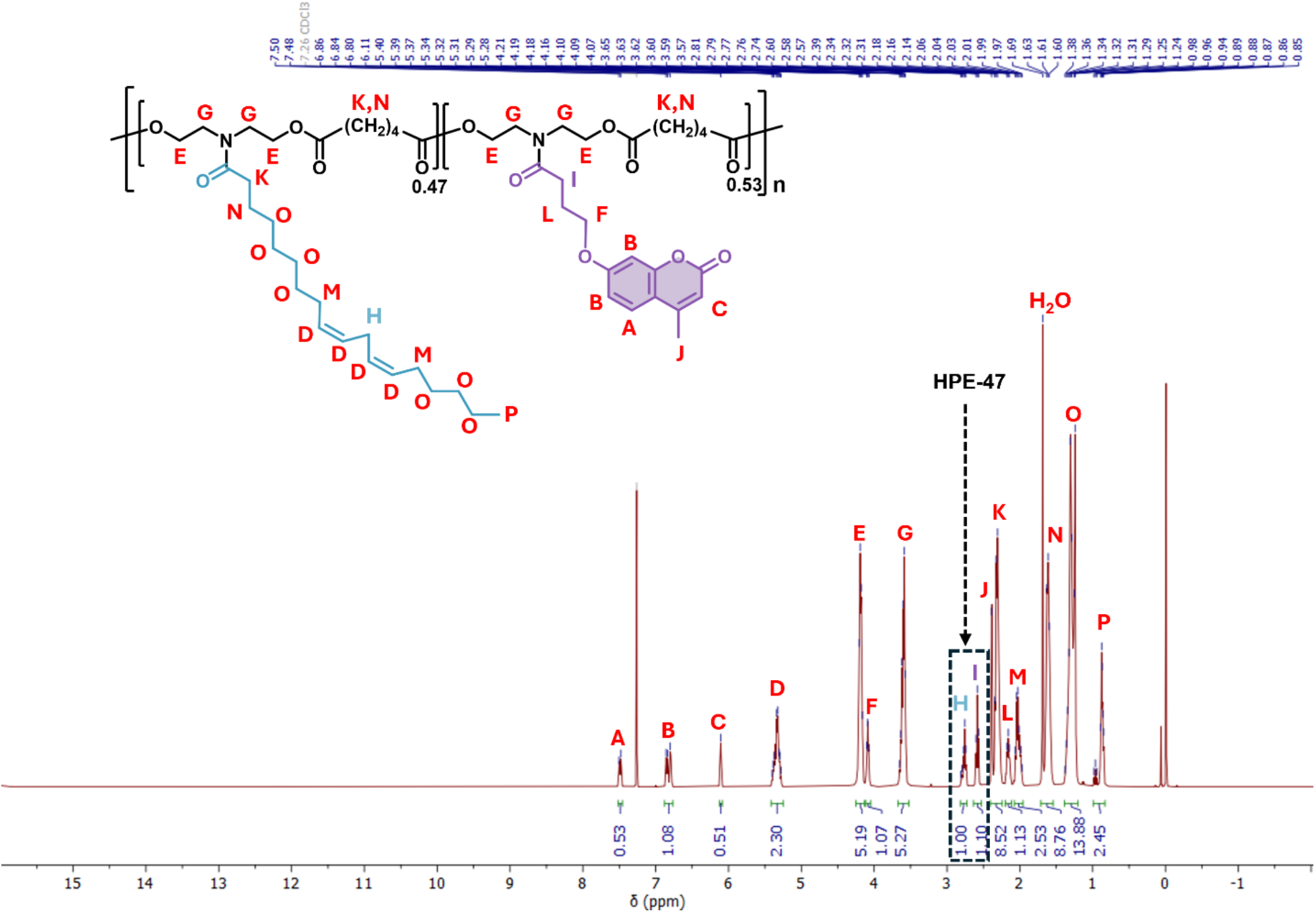
^1^H NMR spectrum of HPE-47 in CDCl_3_.

## 4 : Polymer Characterization Results

### 4.1 Glass transition temperature (T_g_)

Thermal analysis was performed under a nitrogen flow (50 mL min^-1^). Samples underwent two cooling-heating-cooling cycles from − 75 to 120 °C at a rate of 5 °C min^-1^ with 5 min isothermal holds at each temperature extreme (Figure S9).

**Figure S9.**
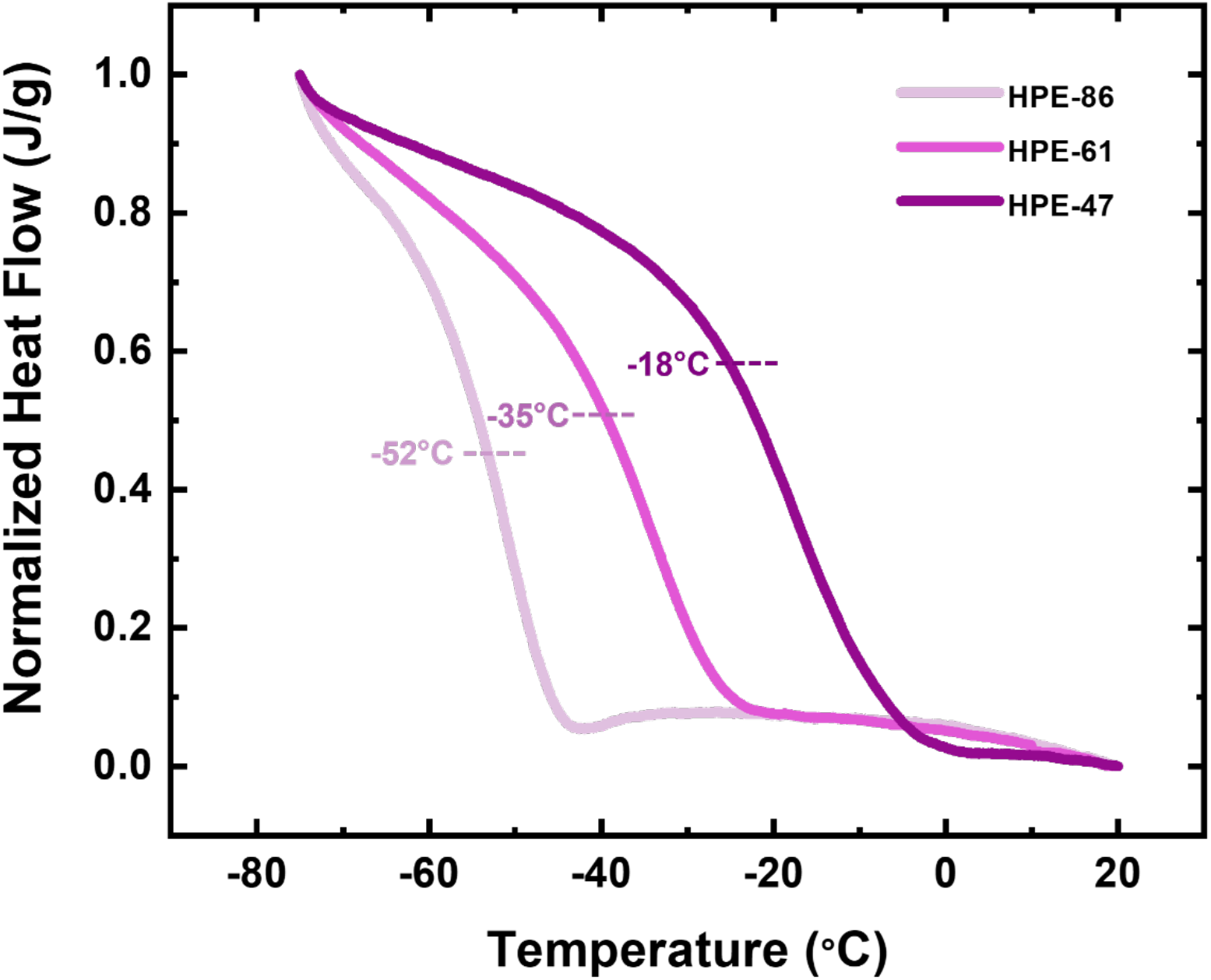
DSC plot summarizing *T*_g_ for HPE-86, HPE-61, and HPE-47. The *T*_g_ increased with decreasing mOil:mPho ratio, consistent with the expected behavior of incorporating coumarin-containing monomers into the polymer.

**Figure S10.**
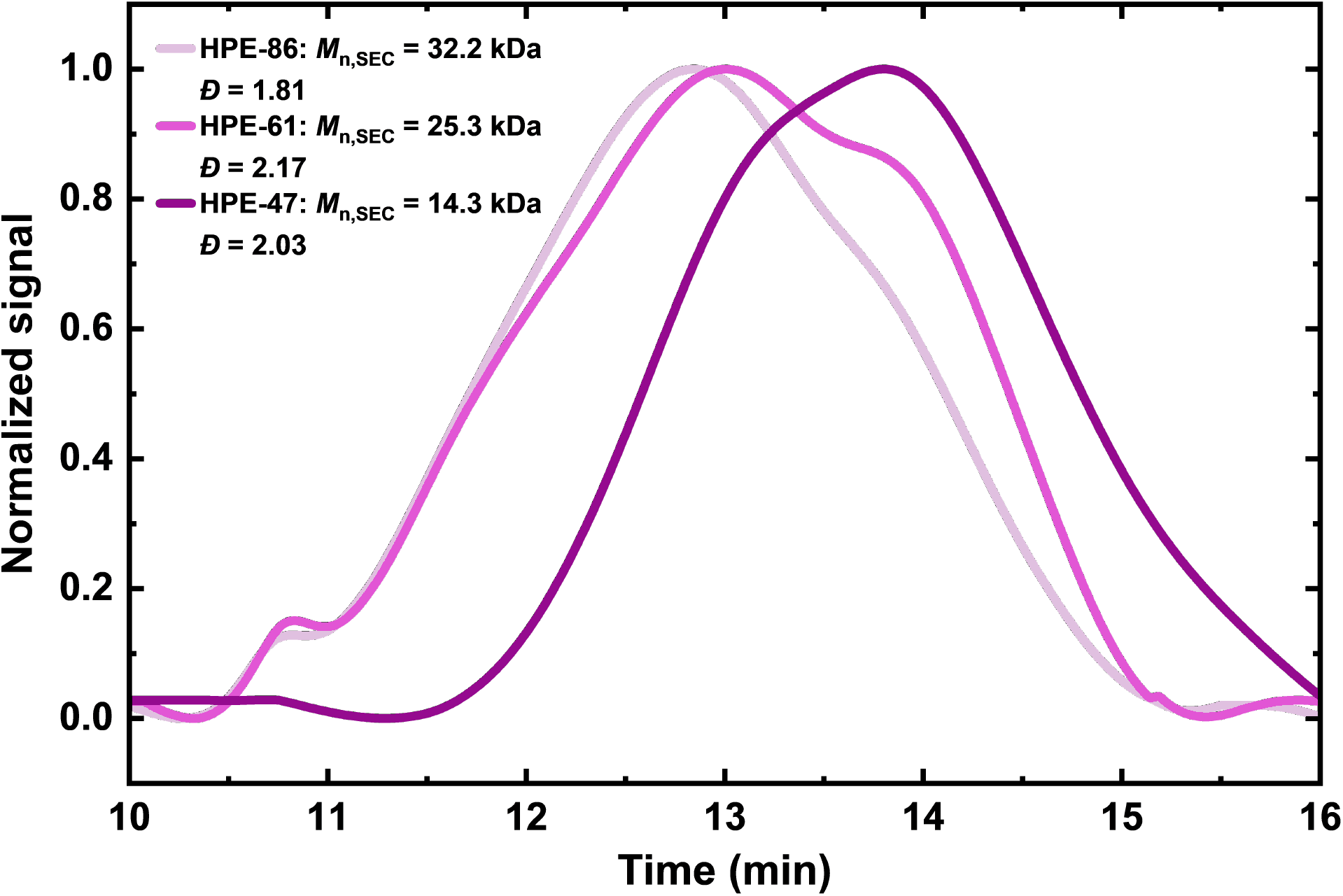
Normalized SEC traces for HPE-86, HPE-61, and HPE-47. THF was used as the eluent and polystyrene as the standard. The legend shows *M*_n,SEC_ (number-average molecular weight) and *Ð* determined from SEC.

### 4.2 Rheology of HPE and PDMS

Rheological measurements were performed on a TA Instruments Discovery HR-2 rheometer using a 20 mm parallel-plate geometry at 25 °C.

Steady-state flow measurements were conducted over a shear rate range of 0.01–100 s^-1^. All HPE exhibited Newtonian behavior at low shear rates, with a transition to non-Newtonian behavior occurring at approximately 10 s^-1^ (Figure S11). Zero-shear viscosity values were extracted from the low-shear rate linear regime.

Oscillatory strain sweep measurements were performed from 0.1 to 1000 % strain at a constant angular frequency of 10 rad s^-1^ (Figure S12). The storage (*G’*) and loss (*G’’*) moduli remained constant within the linear viscoelastic region up to 10 % strain, with nonlinear behavior observed above 100 % strain. For crosslinked HPE, the oscillatory strain sweep measurements were performed from 0.1 to 10 % strain at a constant angular frequency of 10 rad s^-1^ (Figure S13). Across all crosslinking times, *G’* and *G’’* moduli remained constant within the linear viscoelastic region up to ∼ 2 % strain. For PDMS, the oscillatory strain sweep measurements were performed from 0.1 to 100 % strain and showed predominantly elastic solid behavior (Figure S14).

Small-amplitude oscillatory shear measurements were conducted for HPE-86 (Figure S15). The samples were subjected to varying crosslinking times (0, 20, 40, and 60 min) with UV light (wavelength, λ ∼ 365 nm, intensity = 7 mW cm^-2^), using new samples for each condition. Measurements were performed at 0.1 % strain (within the linear viscoelastic region) over an angular frequency range of 0.1–250 rad s^-1^.

**Figure S11.**
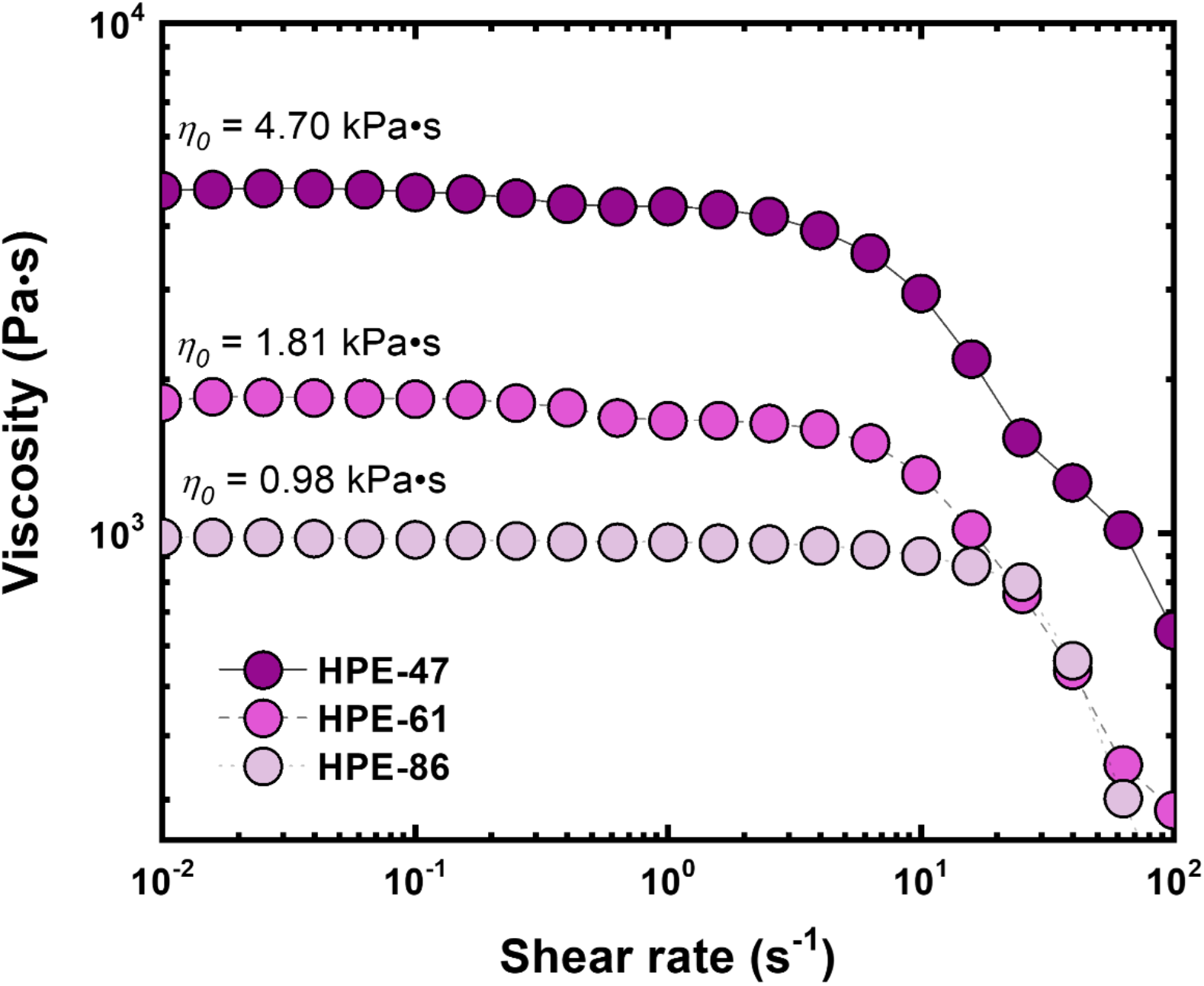
Steady-state flow measurements for HPE-86, HPE-61, and HPE-47 at 25 °C. The zero-shear viscosities (*η*_0_) are shown.

**Figure S12.**
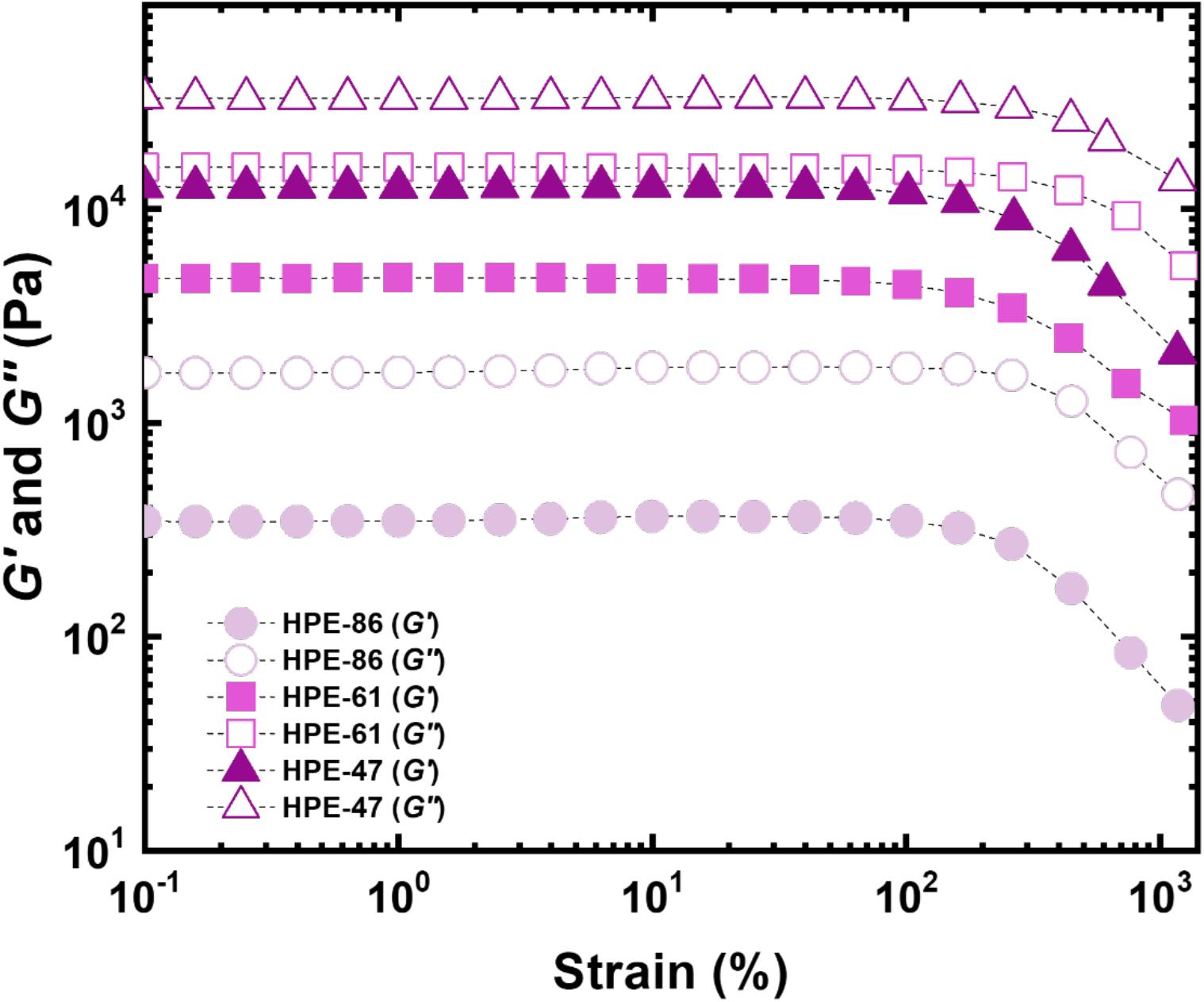
Strain amplitude sweep plot at 10 rad s^-1^ for HPE-86, HPE-61, and HPE-47 at 25 °C. *G’* and *G’’* increase with decreasing mOil:mPho ratio.

**Figure S13.**
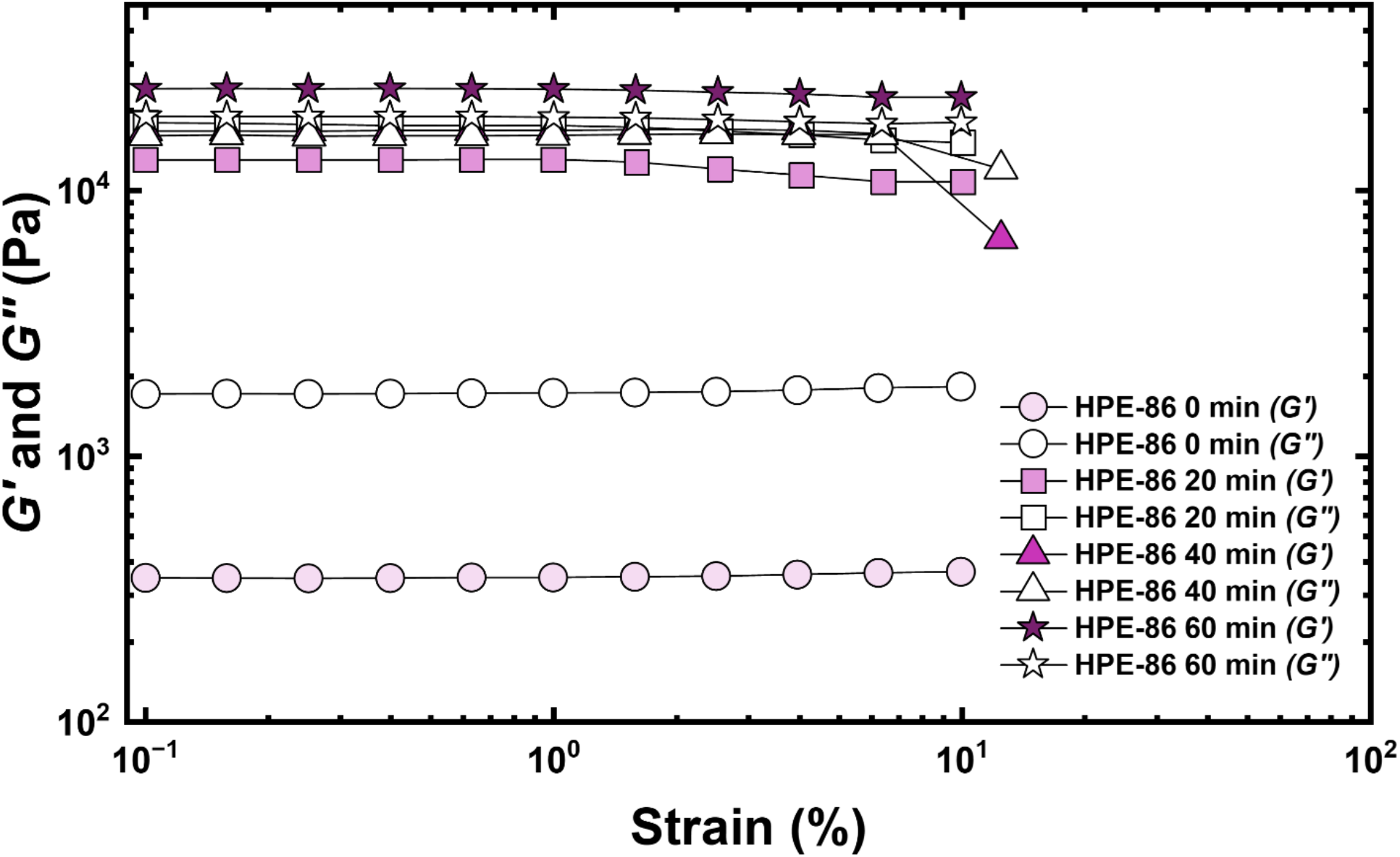
Strain amplitude sweep plot of HPE-86 at 10 rad s^-1^ with varying crosslinking times at 25 °C. The measurements show that crosslinking alters the viscoelastic properties of HPE-86 adhesive. Both the *G’* and *G’’* increased with increasing crosslinking time. Beyond 40 min, *G’* surpassed *G’’*, indicating a transition from viscous-liquid to elastic-solid behavior.

**Figure S14.**
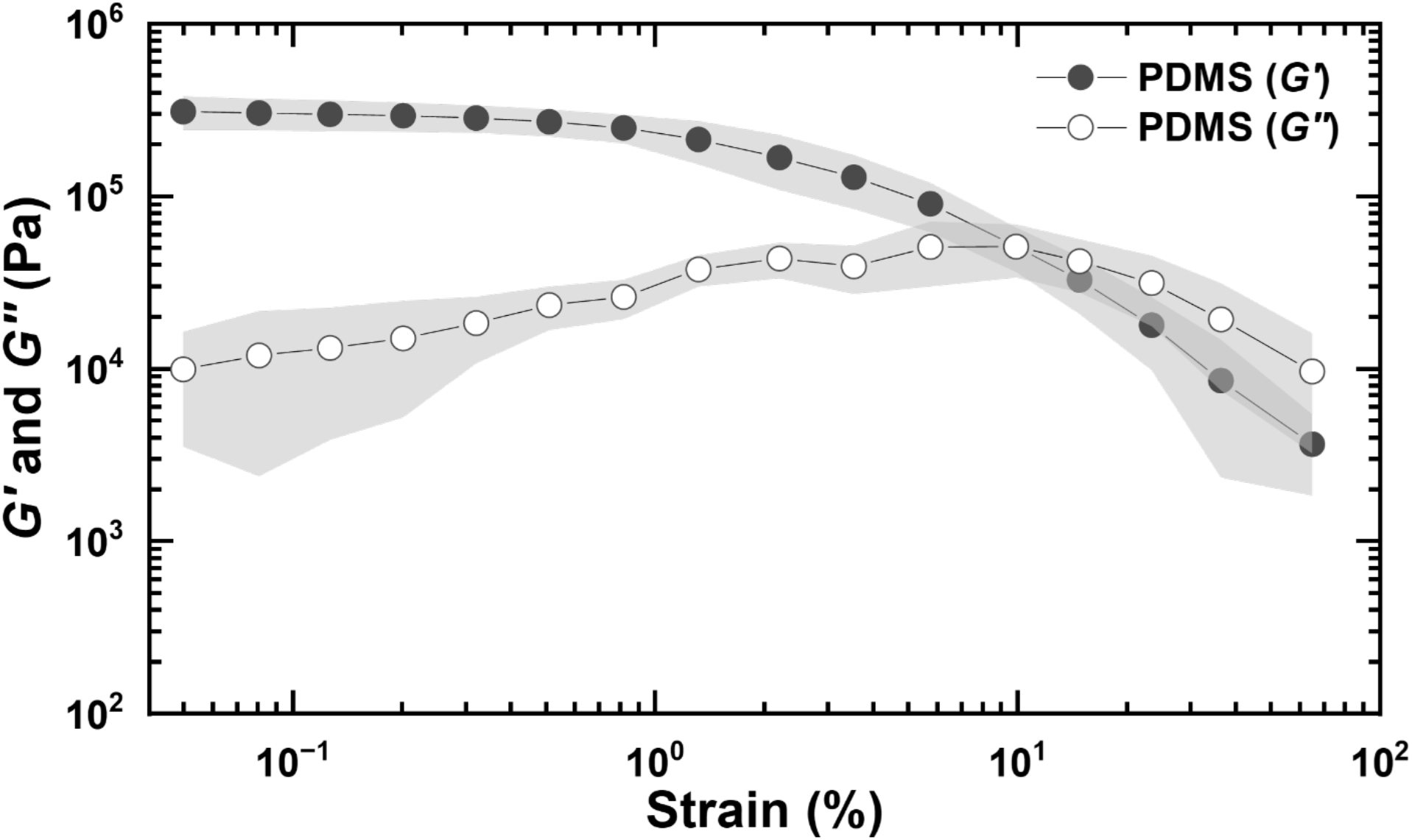
Strain amplitude sweep plot of PDMS at *ω* = 10 rad s^-1^ at 25 °C showing predominantly elastic solid behavior.

**Figure S15.**
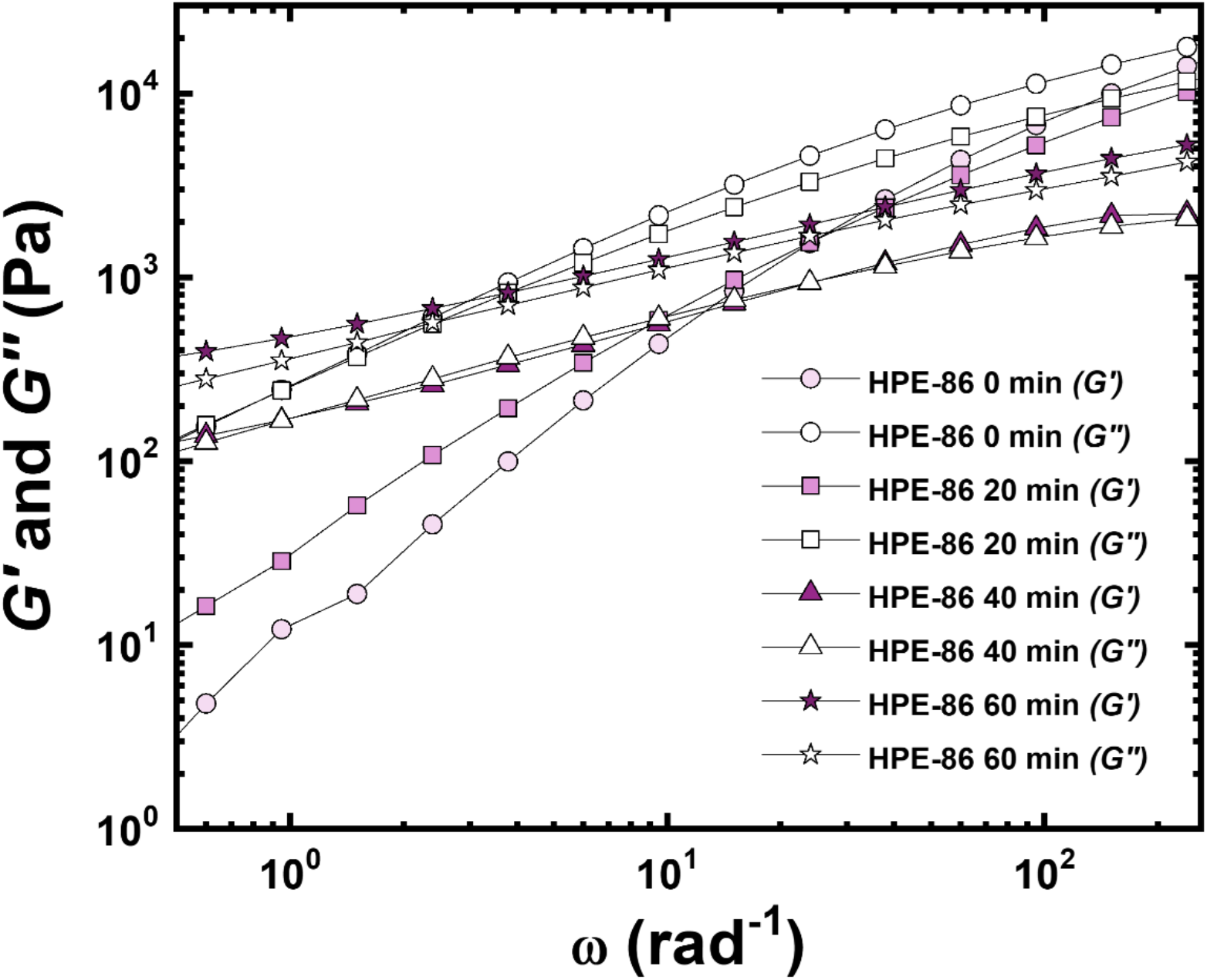
Small-amplitude oscillatory shear measurements of HPE-86 at varying crosslinking times. The experiments were performed at 25 °C. Both *G’* and *G’’* increased with increasing crosslinking, indicating a shift towards more solid-like material behavior.

### 4.3 Pull-off force measurement of HPE on glass under dry and wet conditions

**Figure S16.**
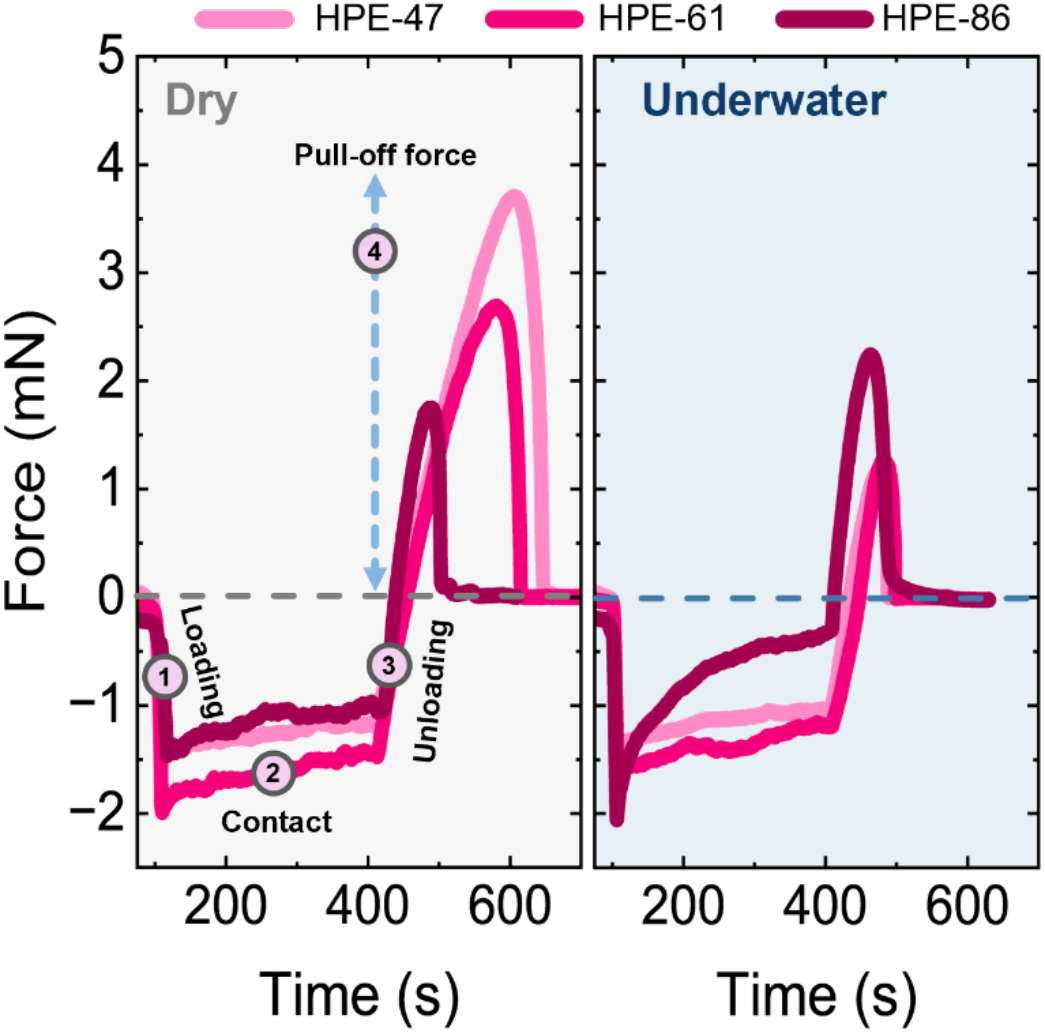
Representative force profiles of HPE samples on a glass substrate in dry and underwater conditions.

## 5 : Biofilm Studies

### 5.1 Biofilm peeling experiments

The general experimental workflow is shown in Scheme S4. Biofilm culture was adapted from Vishwakarma et al.^1^

Bacterial cultures were first grown in LB medium under shaking at 37 °C for 24 h. Cultures were then centrifuged at 3000 rpm for 5 min, the supernatant was discarded, and the cell pellet was inoculated and resuspended in M9 minimal medium. The suspension was grown until reaching OD_600_ ∼ 0.5 and subsequently diluted in M9 medium to ∼10^6^ CFU mL^-1^. Approximately 800 µL of the diluted culture was added to each well of a 24-well glass-bottom plate.

For biofilms grown on SAM-coated surfaces, glass coverslips with self-assembled monolayers (SAMs) were placed in the wells prior to inoculation. In all cases, the plates were sealed with parafilm and incubated statically at 37 °C for 48 h.

Following incubation, the supernatant was removed after centrifugation (3000 rpm, 5 min). To each well, 800 µL of fresh M9 medium was added. Biofilms were stained with DMAO for 30 min in the dark to a final DMAO concentration of 0.19 µM. Excess dye and planktonic cells were removed using a pipette, and 0.85 % NaCl solution was added prior to imaging, per the manufacturer’s protocol.

The confocal microscopy imaging of the peeling process was recorded using 488 nm and 561 nm laser excitation. Before and after Z-stack images were acquired using identical settings (5 % laser power, 40 µm total thickness, 200 nm step size, exposure time). Different samples used varied exposure times. Background images containing only NaCl solution were acquired using the corresponding imaging settings and used for background subtraction during image quantification.

The peeling experiments were performed using three probe types: (i) PDMS, (ii) surface-oxidized PDMS (ox-PDMS), and (iii) HPE-coated PDMS. ox-PDMS was prepared by treating PDMS with a corona torch for 30 s. For HPE-coated samples, PDMS was first corona-treated for 30 s to promote wetting, followed by dip-coating in the HPE solution (4.3 wt% in toluene) for 30 s. This coating was dried under ambient conditions for 2 h, followed by vacuum drying for 24 h to remove residual solvent.

The probes were brought into contact with the biofilm at 1000 nm s^-1^ until the target contact load (2, 5, or 10 mN) was reached (Figure S17 and Figure 1B). This load was maintained for 5 min for equilibration, followed by probe retraction at 30 nm s^-1^. Biofilm removal was quantified using CellProfiler by comparing fluorescence intensity before and after peeling at the biofilm-adhesive contact area.^6^ The image analysis was restricted to the contact region segmented manually, defined as the area showing disruption in the DMAO-emission channel (PDMS or ox-PDMS) or rhodamine channel (HPE). Cohesive failure of the HPE-86 0 min with *S. aureus* was visualized using the rhodamine channel. All images used for biofilm removal quantification were analyzed in their native, unaltered state, without LUT adjustments or other intensity-modifying image processing.

**Scheme S4.**
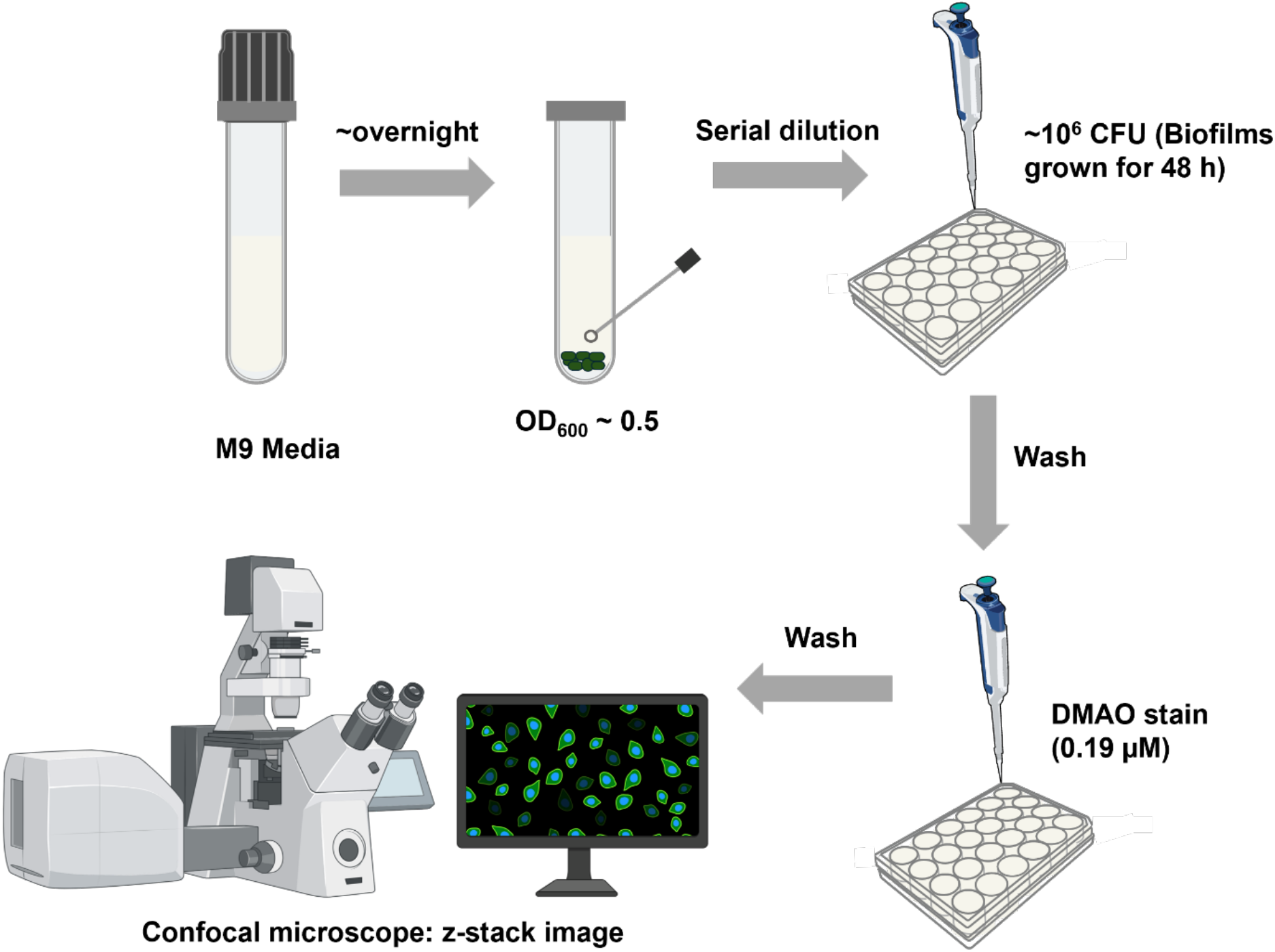
General procedure for the biofilm preparation and quantification using a confocal microscope.

### 5.2 Growth of self-assembled monolayers (SAMs) on glass coverslips

SAM preparation followed a previously reported protocol.^1^ Glass coverslips were cleaned by sonication in solvent baths (toluene → acetone → EtOH) for 10 min. Coverslips were then treated with acid piranha solution, rinsed thoroughly with ultrapure water, and dried with N_2_. The coverslips were further oxidized with corona treatment before immersion in a 0.1 % (v/v) solution of PTS or OTS in anhydrous toluene under continuous N_2_ bubbling. After overnight reaction, the coverslips with SAMs were sonicated sequentially in toluene → acetone → EtOH, dried with N_2_, and annealed at 120 °C overnight before use. The presence of SAMs on the coverslip surfaces was confirmed using water contact angle measurements.

### 5.3 Preparation of hemispherical PDMS elastomer lenses and HPE coating

PDMS elastomer lenses were prepared using Sylgard 184 (10:1 base-to-curing agent ratio) per the manufacturer’s protocol.^7^ The mixture was stirred for 5 min, and degassed under vacuum for 30 min to remove air bubbles. After degassing, the mixture was preheated for 5 min at 80 °C and dispensed (∼ 50 µL) onto a glass substrate precoated with 1H,1H,2H,2H-perfluorodecyltrichlorosilane underwater.^8^ PDMS droplets were cured underwater at 70 °C for 24h. After curing, PDMS lenses (diameter ∼ 2.5 mm) were recovered and stored in a vacuum-sealed desiccator.

Prior to coating with HPE, PDMS lenses were treated with a corona torch for 30 s. A 4.3 wt% HPE solution in toluene was applied via dip-coating for 30 s while maintaining the lens in an inverted orientation to prevent flow-driven accumulation. Toluene was evaporated at room temperature for 2 h, followed by drying under vacuum for 24 h. Samples were stored in a desiccator protected from light to avoid premature crosslinking.

**Figure S17.**
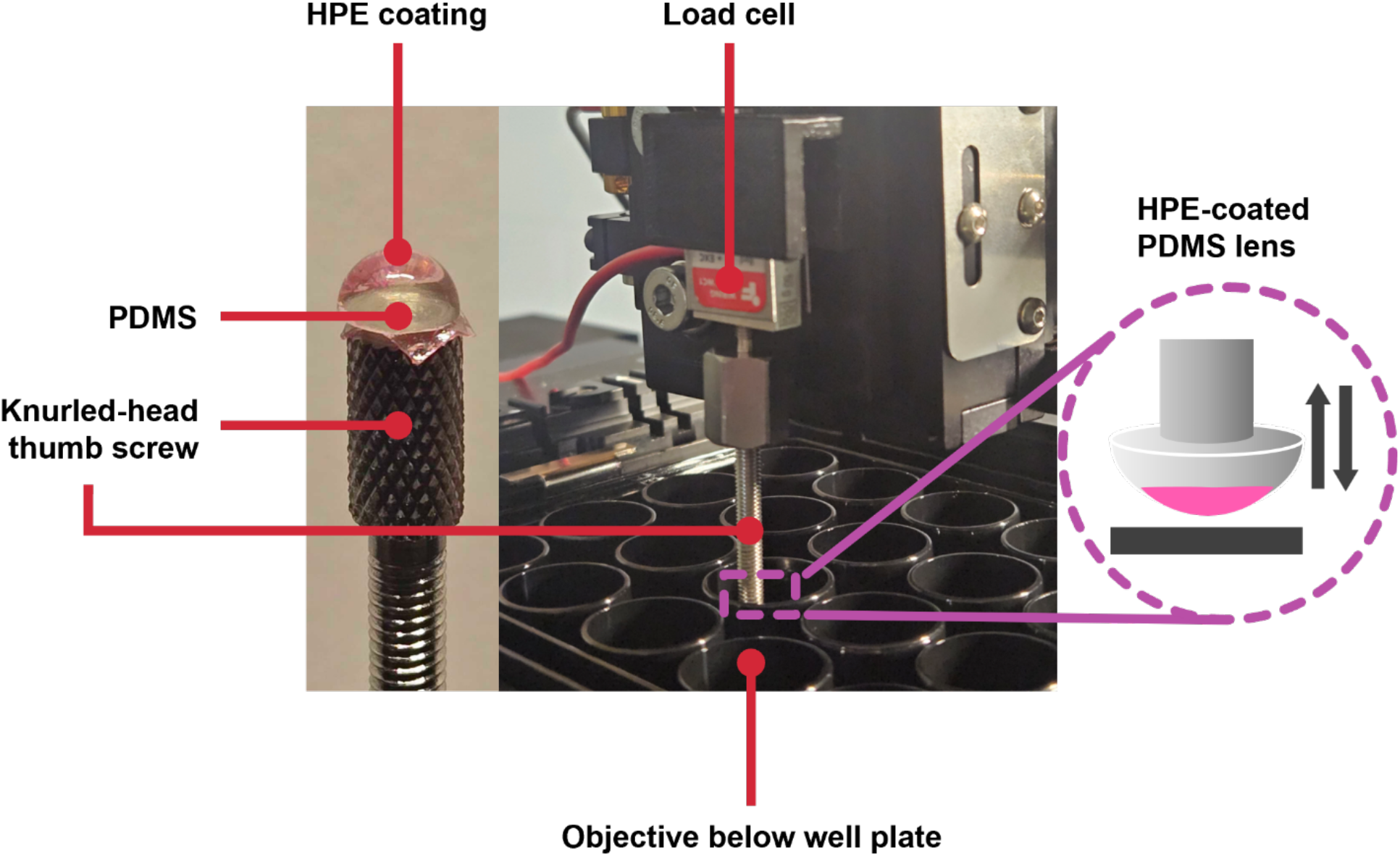
Experimental set-up for underwater biofilm peeling and simultaneous force measurements. HPE-tagged with rhodamine B is coated onto the PDMS. The PDMS is attached to a screw that is connected to a load cell, which is mounted on a piezo actuator to control the peeling speed. A confocal microscope positioned beneath the sample enables real-time imaging and quantification of biofilm removal during peeling.

### 5.4 Biofilm rheology and determination of mechanical mismatch parameter (α)

Biofilms were aseptically inoculated onto tryptic soy agar plates and incubated for 48 h at 37 °C. The visible biofilm colonies (diameter ∼ 0.50 to 1.00 mm) were individually collected and transferred to a 24-well plate.^9^ Plates were centrifuged at 1000 rpm for 5 min to distribute the biofilm evenly in the wells before rheological testing.

Rheology was performed on a TA Instruments Discovery HR-2 rheometer with 8 mm parallel plate geometry at 25 °C (Figure S18). Strain sweeps were performed at a constant angular frequency of 1 rad s^-1^ over a range of 0.01 to 100 % strain (Figure S19). Three replicates were performed for this analysis.

The complex modulus (*G\**) is a measure of a material’s resistance to deformation under an oscillating shear strain, wherein the contributions coming from both elastic (*G’*) and viscous (*G’’*) responses are combined as follows:

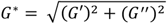

A higher *G\** indicates a stiffer material that can resist deformation more than a material with a lower *G\** value.

The mechanical mismatch (*α_BA_*) between the biofilm and adhesive was used to evaluate the contrasting stiffness between these two materials. This parameter is calculated from their respective *G\** values as shown in the equation (Table S3).

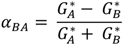

*G_A_*^∗^ and *G_B_*^∗^ are complex moduli of the adhesive and biofilm, respectively, measured within the linear regime at 0.1 % strain. In this study, the values are summarized in Tables S1 and S2.

**Table S1.**
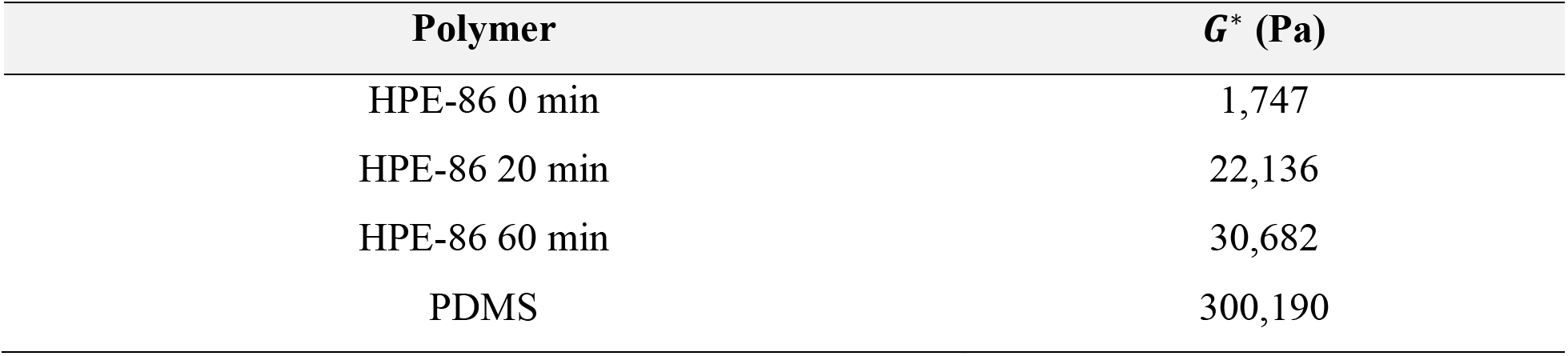
*G_A_*^∗^ values at 0.1 % strain for HPE-86 at varying crosslinking times and for PDMS.

| **Polymer** | ***G***<sup>\*</sup> **(Pa)** |
|---|---|
| HPE-86 0 min | 1,747 |
| HPE-86 20 min | 22,136 |
| HPE-86 60 min | 30,682 |
| PDMS | 300,190 |

**Table S2.** *G_B_*^∗^ values at 0.1 % strain for the biofilms.

| **Biofilm** | ***G***<sup>\*</sup> **(Pa)** |
|---|---|
| *E. coli* | 13,240 |
| *S. aureus* | 10,210 |
| *P. aeruginosa* | 11,370 |

**Table S3.** *α_BA_* values of the polymers and biofilms.

| **Adhesive** | **α<sub>BA</sub> (*E. coli*)** | **α<sub>BA</sub> (*S. aureus*)** | **α<sub>BA</sub> (*P. aeruginosa*)** |
|---|---|---|---|
| HPE-86 0 min | -0.77 | -0.71 | -0.73 |
| HPE-86 20 min | 0.25 | 0.37 | 0.32 |
| HPE-86 60 min | 0.40 | 0.50 | 0.46 |
| PDMS | 0.92 | 0.93 | 0.93 |

**Figure S18.**
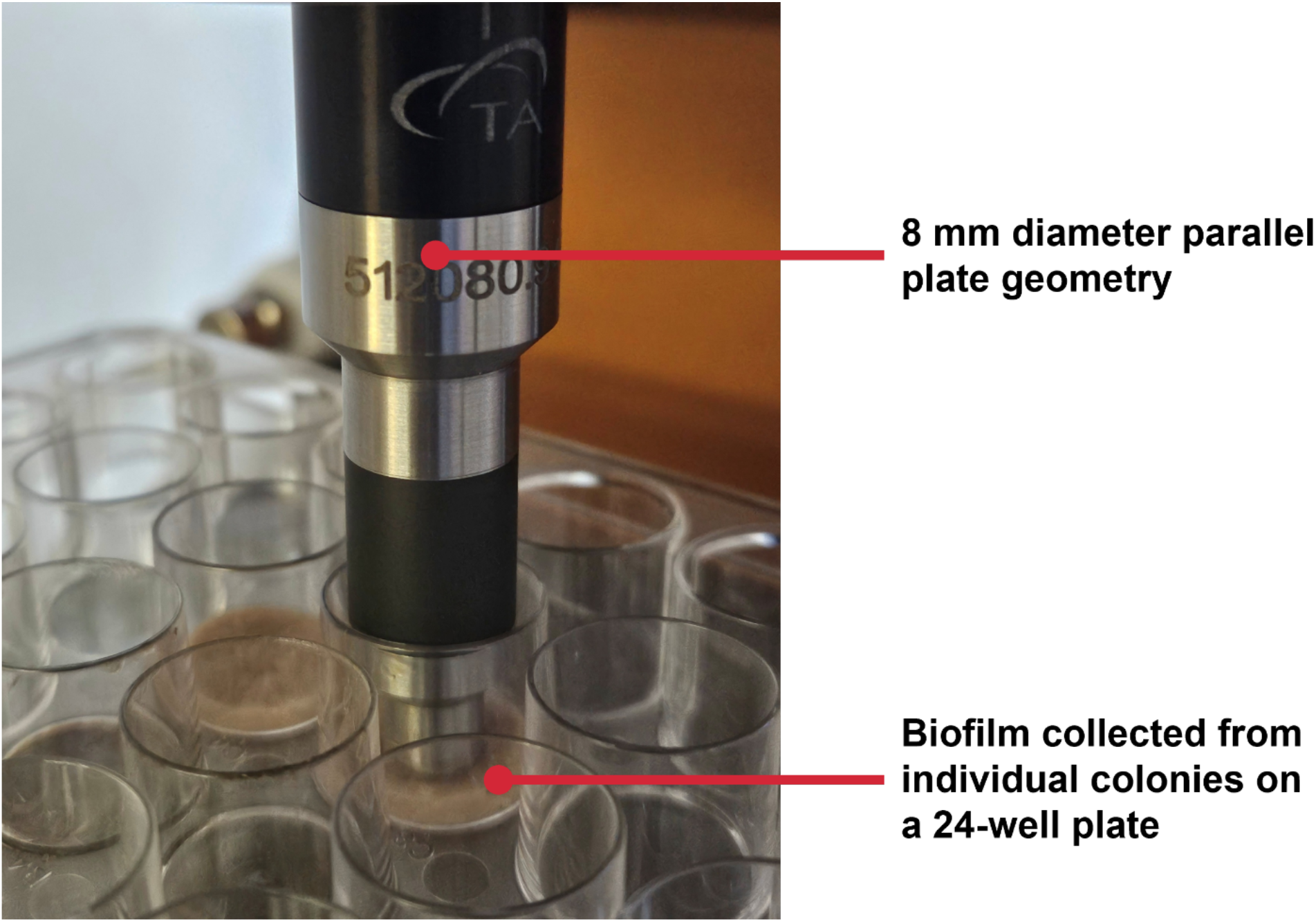
Biofilm rheology set-up showing biofilms that were individually collected and centrifuged on a 24-well plate. 8 mm parallel plate geometry was used in the strain sweep experiments.

**Figure S19.**
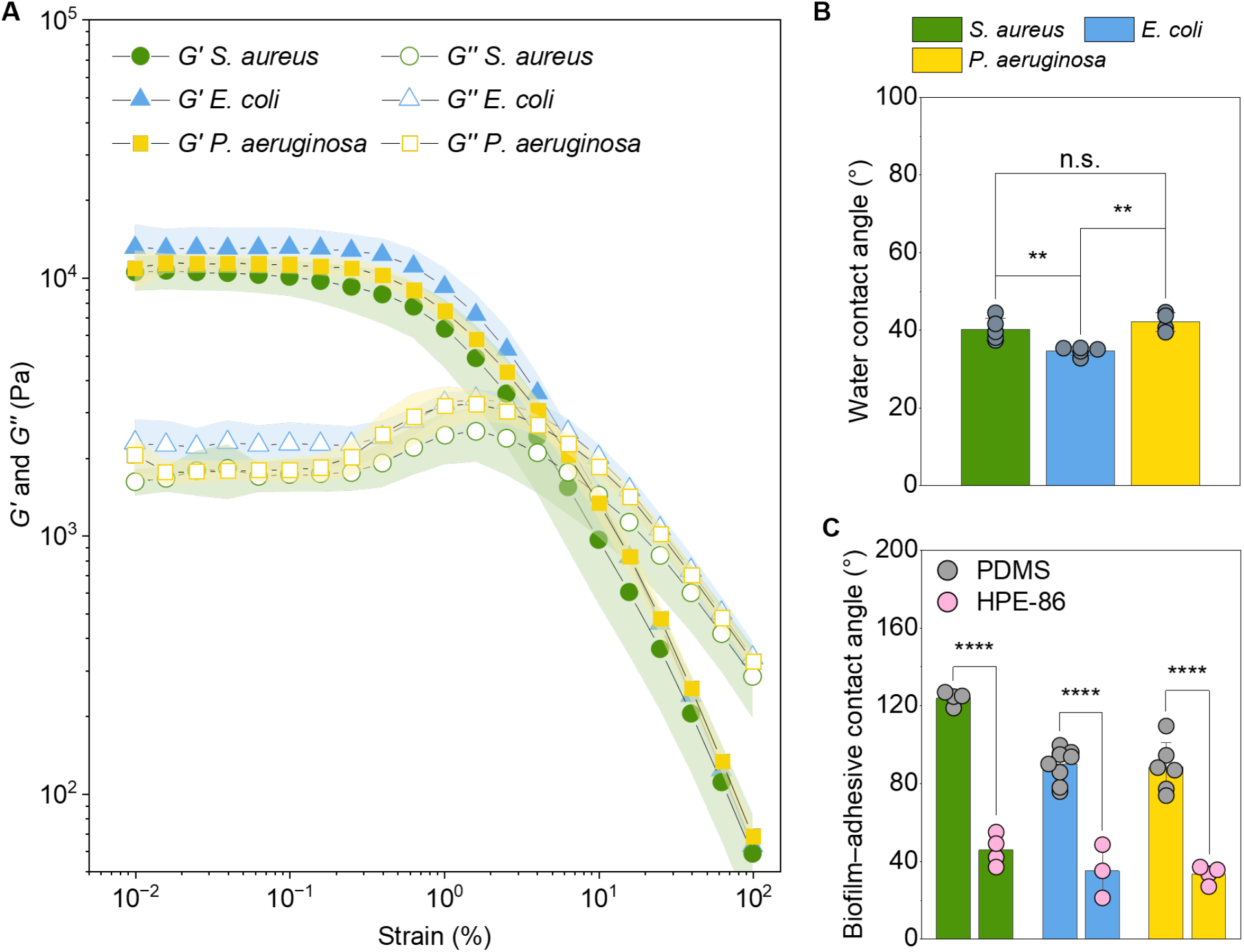
(A) Strain amplitude sweep of *E. coli*, *S. aureus*, and *P. aeruginosa* biofilms from 0.01 to 100 % strain at 25 °C. (B) Water contact angle of biofilms. Bar height represents the mean, error bars show standard deviation, and the circular symbols show individual replicates. Mean comparisons were performed using one-way ANOVA, and the significant differences across individual conditions were determined using Tukey’s test. (n.s. = not significant, \**p* < 0.05, \*\**p* < 0.01, \*\*\**p* <0.001, \*\*\*\**p* < 0.0001). (C) Contact angle of HPE-86 (pink symbol) and PDMS (gray symbol) on biofilms underwater.

## 6 : Contact Angle Measurements and Determining Thermodynamic Work of Adhesion (*W*ad)

### 6.1 Water contact angle experiments on polymer surfaces

The static water contact angles were measured on three surface types: PDMS, ox-PDMS, and HPE-86-coated PDMS. ox-PDMS and HPE-86-coated PDMS were prepared as outlined in section 5.3.

Contact angle measurements were performed at room temperature (Table S4). A 10 µL droplet of deionized water was deposited onto the surface and imaged immediately upon contact. Contact angles were determined using the Contact Angle plugin in ImageJ. The left and right angles were measured and averaged. At least three measurements were performed for each sample type.

**Table S4.** Water contact angle of the adhesives.

| **Adhesive sample** | **Water contact angle (°)** |
|---|---|
| PDMS | 109.9 ± 1.5 |
| ox-PDMS | 26.0 ± 6.3 |
| HPE-86 0 min | 109.6 ± 1.8 |

### 6.2 Water contact angle experiments on biofilms

Static water contact angles were measured on intact biofilms grown on tryptic soy agar plates (Figure S19B and Table S5). Bacteria were aseptically streaked onto tryptic soy agar plates and incubated at 37 °C for 48 h. After incubation, agar sections with biofilm were excised using a sterile blade to minimize disruption of the biofilm surface and preserve its native surface properties.

A 10 µL droplet of deionized water was deposited onto the biofilm surface, and contact angles were measured at room temperature immediately upon contact. Contact angles were determined using the Contact Angle plugin in ImageJ. At least three measurements were performed for each species.

**Table S5.**
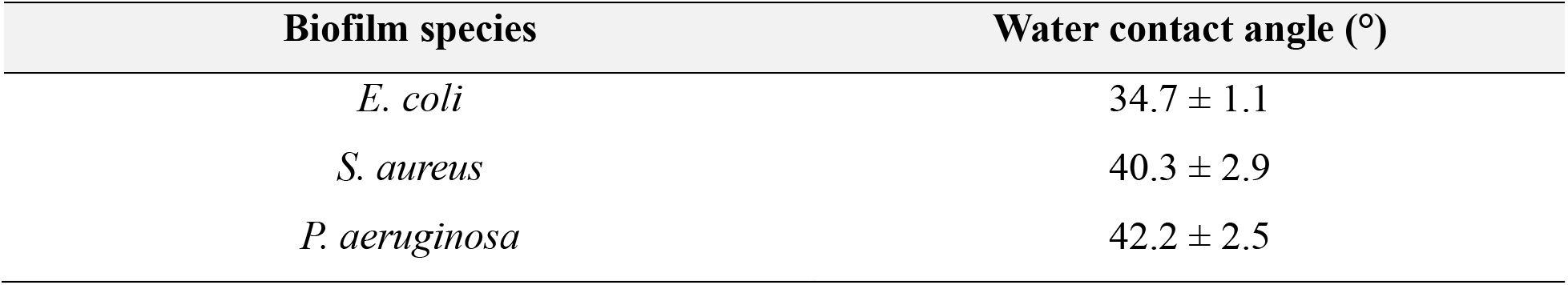
Water contact angle of biofilms.

### 6.3 Underwater contact angle of adhesives on biofilms

Bacteria were grown in LB medium at 37 °C to 0.50 OD_600_, diluted in fresh LB medium to ∼10^6^ CFU mL^-1^, and 800 µL of the suspension was added to each well of a 24-well plate containing glass coverslips. Plates were incubated statically at 37 °C for 24 h. After incubation, the supernatant was removed by pipetting, and the coverslips were rinsed twice with deionized water to remove planktonic bacteria.

Biofilm-coated coverslips were removed from the wells and immediately submerged in deionized water in a glass cell. For PDMS, a 10 µL drop of Sylgard 184 Part A was dispensed onto the biofilm surface using a pipette. Because the PDMS contact angle did not change significantly over time, the angle recorded during initial contact was reported (Figure S19C and Table S6).

For biofilm–HPE-86 contact angle measurements, the HPE was backloaded into a syringe and preheated to reduce its viscosity before ∼ 10 µL was deposited onto the biofilm surface. The water temperature was monitored as the HPE droplet cooled, and the contact angle was recorded once the temperature reached 25 °C (Figure S19C and Table S6). This temperature was selected because the contact angle decreased as the droplet cooled and spread. Contact angles were determined using the Contact Angle plugin in ImageJ. At least three measurements were conducted.

**Table S6.** Polymer contact angle on biofilms.

| **Biofilm samples** | **Contact angle of PDMS underwater (°)** | **Contact angle of HPE-86 underwater (°)** |
|---|---|---|
| *E. coli* | 89.9 ± 8.3 | 35.0 ± 13.7 |
| *S. aureus* | 123.8 ± 3.5 | 45.9 ± 7.9 |
| *P. aeruginosa* | 88.4 ± 12.8 | 33.3 ± 4.4 |

### 6.4 Captive bubble contact angle measurements and the determination of adhesive-water interfacial tension

Adhesive-coated substrates were mounted face down in a glass cell filled with deionized water at room temperature. For HPE-coated substrates, 10 µL of HPE solution (4.3 wt% in toluene) was deposited onto a silicon wafer and spin-coated at the maximum speed of the spin coater for 20 min. Coated wafers were dried under vacuum for 24 h to remove residual solvent.

An air bubble (∼10 µL) was formed using a syringe fitted with a 90° L-shaped needle and released beneath the substrate (Figure S20). Contact angles were measured immediately upon bubble contact (Table S7). The reported angles were measured underwater. At least three measurements were conducted.

The adhesive-water interfacial tension (*γ_AW_*) was calculated from the captive bubble contact angle using the Young’s equation, where *γ_Av_* is the adhesive-air interfacial tension, *γ*_W*v*_ is the water-air interfacial tension, and *θ_Av_* is the contact angle from the captive bubble experiment (Table S7). For HPE-86, a value of *γ_Av_* = 20 mN/m was used for the calculation.^10^

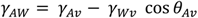

**Table S7.**
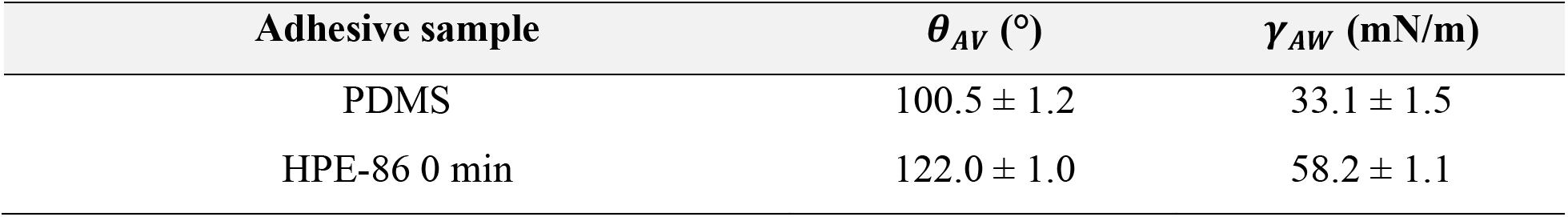
Captive bubble contact angle results and adhesive–water interfacial tension.

| **Adhesive sample** | ***θ<sub>AV</sub>*** **(°)** | ***γ<sub>AW</sub>*** **(mN/m)** |
|---|---|---|
| PDMS | 100.5 ± 1.2 | 33.1 ± 1.5 |
| HPE-86 0 min | 122.0 ± 1.0 | 58.2 ± 1.1 |

**Figure S20.**
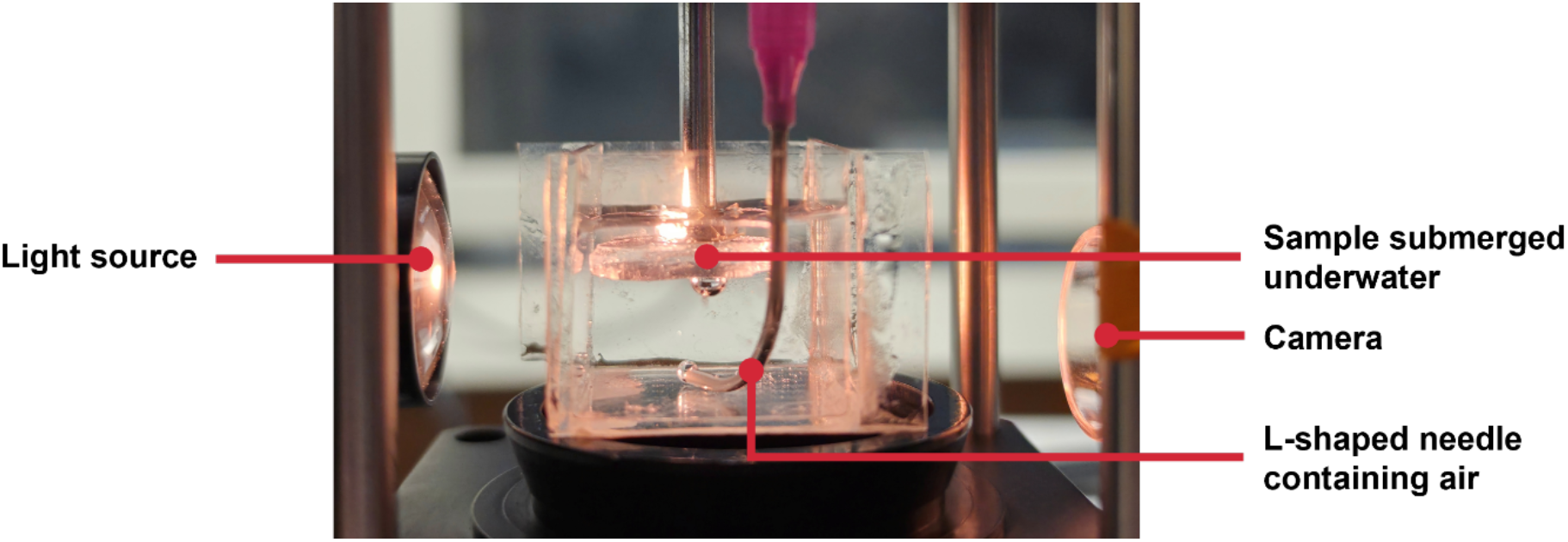
Captive bubble set-up: Approximately 10 µL air bubble was placed against an inverted sample submerged in water. The sample surface consisted of either PDMS or HPE-86-coated silicon wafer. Bare PDMS was used in the set-up shown. The underwater contact angle of the bubble (*θ_Av_*) on the PDMS surface was measured to calculate the adhesive-water interfacial tension (*γ_AW_*).

### 6.5 Determination of biofilm–adhesive thermodynamic work of adhesion

The work of adhesion (*W*_ad_) between each polymer and biofilm pair in the aqueous environment was calculated from the underwater biofilm–adhesive contact angle using the Young-Dupré and Young equations (Table S8). Here, *γ_AW_* is the adhesive-water interfacial tension, while *θ_BA_* is the biofilm–adhesive contact angle measured underwater.

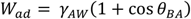

**Table S8.**
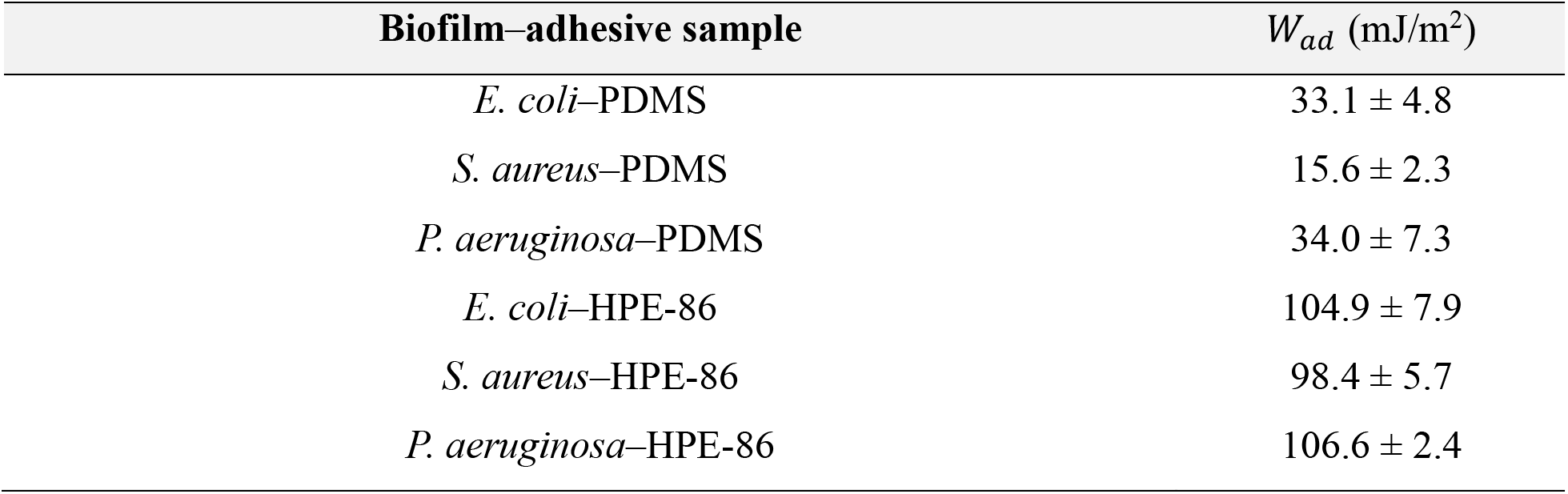
Calculated biofilm–adhesive work of adhesion.

| **Biofilm–adhesive sample** | *W<sub>ad</sub>* (mJ/m<sup>2</sup>) |
|---|---|
| *E. coli*–PDMS | 33.1 ± 4.8 |
| *S. aureus*–PDMS | 15.6 ± 2.3 |
| *P. aeruginosa*–PDMS | 34.0 ± 7.3 |
| *E. coli*–HPE-86 | 104.9 ± 7.9 |
| *S. aureus*–HPE-86 | 98.4 ± 5.7 |
| *P. aeruginosa*–HPE-86 | 106.6 ± 2.4 |

## 7 : Additional Data

**Figure S21.**
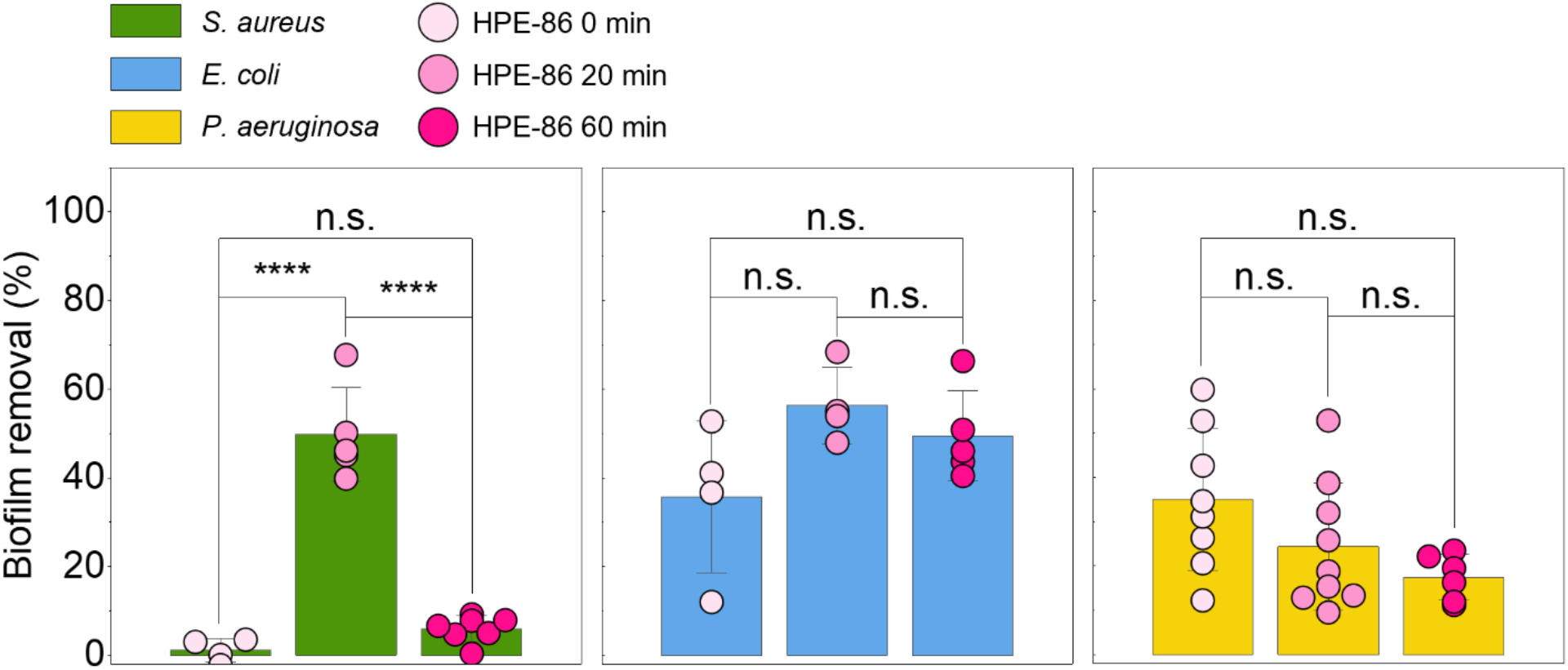
Biofilm removal of HPE-86 0 min, 20 min, and 60 min for *S. aureus*, *E. coli*, and *P. aeruginosa*. Mean comparisons were performed using one-way ANOVA, and the significant differences across individual conditions were determined using Tukey’s test (n.s. = not significant, *p < 0.05, **p < 0.01, ***p <0.001, ****p <0.0001).

**Figure S22.**
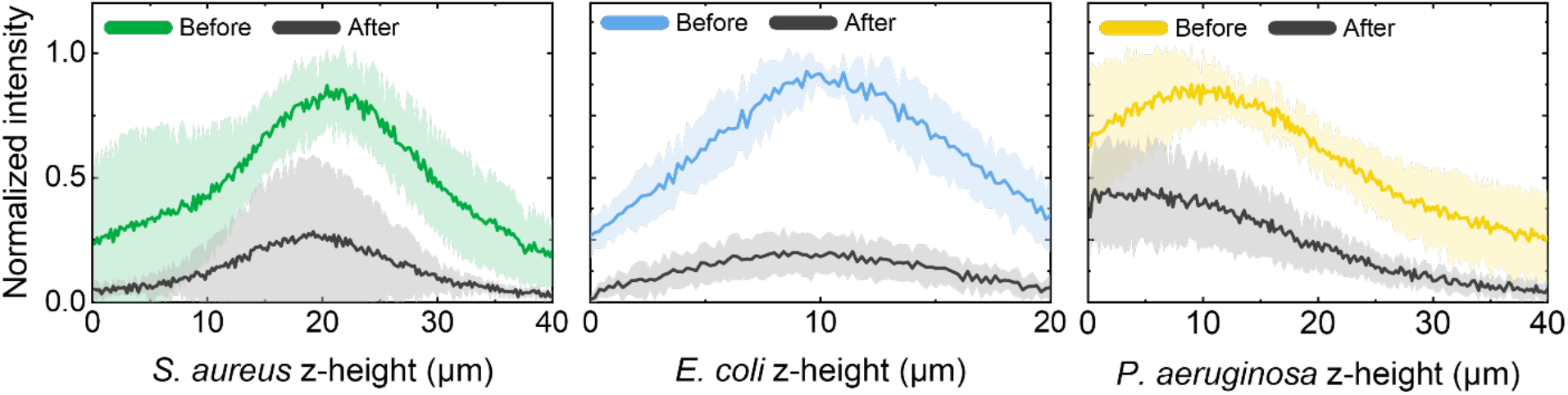
Normalized fluorescence intensity before and after peeling as a function of depth for the *S. aureus*, *E. coli*, and *P. aeruginosa* biofilms peeled using HPE-86 20 min. The analysis is restricted to the contact region at a contact load of 2 mN. The peeling occurs at all Z-heights for *S. aureus* and *E. coli*, while *P. aeruginosa* profiles are peeled from the top region of the biofilm (higher Z-height), indicating cohesive failure of *P. aeruginosa* biofilms during adhesive peeling.

